# A flexible modeling and inference framework for estimating variant effect sizes from GWAS summary statistics

**DOI:** 10.1101/2022.04.18.488696

**Authors:** Jeffrey P. Spence, Nasa Sinnott-Armstrong, Themistocles L. Assimes, Jonathan K. Pritchard

## Abstract

Genome-wide association studies (GWAS) have highlighted that almost any trait is affected by many variants of relatively small effect. On one hand this presents a challenge for inferring the effect of any single variant as the signal-to-noise ratio is high for variants of small effect. This challenge is compounded when combining information across many variants in polygenic scores for predicting trait values. On the other hand, the large number of contributing variants provides an opportunity to learn about the average behavior of variants encoded in the distribution of variant effect sizes. Many approaches have looked at aspects of this problem, but no method has unified the inference of the effects of individual variants with the inference of the distribution of effect sizes while requiring only GWAS summary statistics and properly accounting for linkage disequilibrium between variants. Here we present a flexible, unifying framework that combines information across variants to infer a distribution of effect sizes and uses this distribution to improve the estimation of the effects of individual variants. We also develop a variational inference (VI) scheme to perform efficient inference under this framework. We show this framework is useful by constructing polygenic scores (PGSs) that outperform the state-of-the-art. Our modeling framework easily extends to jointly inferring effect sizes across multiple cohorts, where we show that building PGSs using additional cohorts of differing ancestries improves predictive accuracy and portability. We also investigate the inferred distributions of effect sizes across many traits and find that these distributions have effect sizes ranging over multiple orders of magnitude, in contrast to the assumptions implicit in many commonly-used statistical genetics methods.

## 1 Introduction

The central problem in statistical genetics is understanding the role of genetic variation in shaping observed phenotypic variation. This map from genotype to phenotype can be understood at multiple scales of granularity. At one extreme, elucidating the functional impact of individual variants can provide biological insights into molecular mechanisms or pathways [51] and for phenotypes related to disease status can be used for drug discovery [13, 44]. At the other extreme, we can discover broad features of the genotype to phenotype map without knowing the effect of any particular variant. For example, with polygenic scores (PGSs) we can use genotypes to predict disease risk to implement more cost-effective screening programs without accurately knowing how any single variant is acting [17, 20, 33]. We can also learn summaries of the distribution of effect sizes such as the proportion of phenotypic variance explained by genetic variation (heritability), how polygenic a trait is, or if particular types of genomic regions are enriched for contributions to a trait. Looking across traits or across cohorts, we can analogously consider the joint distribution of effect sizes and infer the extent to which the effect of variants is correlated across those traits or cohorts. Across all these scales of granularity there are interesting questions and important medical applications.

Unfortunately, there are challenging statistical problems at all scales. Correlations between genotypes at nearby loci (linkage disequilibrium; LD) make it difficult to disentangle the effect of a variant at one locus from the effects of correlated variants. Compounding these issues, genome wide association studies (GWAS) often only release the summary statistics of marginal tests of association performed separately for each variant, which do not adjust for the effect of linked variants. As a result, methods must account for LD while only having access to these summary statistics.

When inferring the effects of particular variants there is an additional problem: at present sample sizes, there is no simple way to estimate the effect of a variant while accounting for all other variants. Sample sizes are often in the thousands to the hundreds of thousands, while the number of variants can be in the millions, making tools from classical statistics like multiple regression impossible to apply. In principle, Bayesian methods or regularization methods such as the LASSO [31, 57] or ridge regression [24, 59] can make the original ill-posed problem well-posed. Yet, without a solid understanding of the distribution of effect sizes, choosing the form and amount of regularization can be difficult. For example, LASSO regularization favors sparse solutions where only a small proportion of variants have non-zero effects, and ridge regression favors solutions where no single variant has a large effect. For many complex traits neither of these assumptions is appropriate: there are often variants with relatively large effects, but simultaneously the vast majority of phenotypic variance is explained by many variants of tiny effect [8, 51].

While having a large number of variants can be thought of as a typical “curse of dimensionality” for inferring the effect of any particular variant, in the context of learning about the distribution of effect sizes the large number of variants can also be seen as a blessing. Each variant provides some information about the distribution of variant effect sizes, and so by pooling information across variants we might hope to discover features of this distribution. Using an estimated distribution of effect sizes, one can apply more sensible regularization when inferring the effects of individual variants. This highlights that while we can try to understand the genotype to phenotype map at each resolution or degree of granularity separately, there is information to be gained by considering all scales jointly.

There have been many approaches to interrogate these different aspects of the genotype to phenotype map, but no sufficiently flexible unifying framework has been developed. Many methods have been developed for estimating individual effect sizes, usually within a single genetic locus, especially under the assumption than only one or a small number of variants are causal. This setting is known as fine-mapping, and some of these methods require only GWAS summary statistics [2, 61, 67]. A related line of work looks at multiple traits (often an organism-level phenotype and gene expression) and tries to “colocalize” signals by determining if the same variants can explain observed associations with both traits [25]. Yet, these methods typically do not use information about the overall distribution of effect sizes to inform their predictions. On the other end of the spectrum, there are a number of methods that use either genotype data or summary statistics to estimate features of the distribution of effect sizes without estimating the effects of individual variants. In particular, variance component models and models based on the LD Score Regression framework have been used to estimate heritability, which is related to the variance of the distribution of frequency-scaled effect sizes [10]. These models have also been used to estimate other aspects of this distribution of effect sizes, such as its fourth moment, a measure of how heavy-tailed this distribution is [39]. For multiple traits or cohorts, there are methods that can estimate the correlation of effect sizes [9]. Finally, recent work has looked at estimating the full distribution of contributions to heritability, which is closely related to the distribution of effect sizes [38]. A series of methods that do leverage information in the variants to jointly estimate the effect size distribution and then use that inferred effect size distribution to improve the estimation of individual effect sizes have also been developed, but these require the variants to be independent. This side-steps issues of LD but necessitates throwing away the information contained in linked SNPs [55].

A related line of work predicts phenotype from genotype using so-called “polygenic scores” (PGSs) or “polygenic risk scores”. State-of-the-art approaches use some form of explicit regularization like the LASSO [31], or perform Bayesian inference, where an assumed distribution of effect sizes is used as a prior and acts as a regularizer. These methods typically specify a particular family of priors such a Normal with a point mass at zero [59], mixture of a small number of Normals [30], or a particular scale-mixture of Normals [22], and the user is required to choose a distribution from this family by tuning a hyperparameter using a held-out validation dataset. Recent work has eliminated the need for a validation dataset by placing an additional prior on the hyperparameters and then obtaining a posterior distribution over assumed effect size distributions [22, 65], but this prior then contains hyperparameters which are fixed *a priori*. Furthermore, these methods, with the exception of [65], often make restrictive and unrealistic assumptions about the distribution of effect sizes, such as only having effect sizes from roughly a single order of magnitude [59], or coupling the probability that an effect size is close to zero with the probability that an effect size is large [22]. Currently, only one method in this framework, PRS-CSx [45], models effect sizes across cohorts, and this method implicitly assumes that while the magnitude of effects are similar across cohorts, the genetic correlation across cohorts is zero in contrast to what is seen in real data [9]. As such, while many aspects of the central problem of statistical genetics have been tackled in isolation, no single method combines inference of a sufficiently flexible distribution of effect sizes while also using such information to inform the inference of the effects of individual variants.

Here we present a unified framework that ties an extremely flexible, learnable family of effect size distributions to GWAS summary statistics, allowing for the simultaneous estimation of the effects of individual variants along with the distribution of effect sizes. We extend our method to the case of multiple cohorts, potentially with distinct LD structures, where we can learn the joint distribution of effects across cohorts and use information contained in multiple GWAS to improve estimation of variant effect sizes across all cohorts. Our model possesses several key features: 1) it is flexible, allowing effect sizes to vary across multiple orders of magnitude with varying degrees of genetic correlation across cohorts; 2) it properly accounts for LD while only requiring the use of GWAS summary statistics, making it amenable to use on publicly available data; 3) our model can be fit efficiently using modern tools from variational inference (VI); 4) our model can incorporate prior information, such as genomic annotations or molecular data, by using this information to create site-specific effect size distributions; and 5) our model is easily extendable to model multiple traits instead of multiple cohorts.

As an example of the utility of our model, we focus specifically on the case of building PGSs. PGSs are used to predict an individual’s phenotype or risk of disease and are becoming accurate enough to be clinically useful [27]. Yet, PGSs typically explain far less trait variance than theoretically possible – the narrow-sense heritability – showing that there is still room for improvement. Here, we show that PGSs derived using our framework can be substantially more predictive than the current state-of-the-art method [22, 45].

PGSs typically suffer from poor portability, wherein PGSs built using data from a particular cohort perform far worse when applied to sets of individuals that are genetically distant from the cohort used to build the PGS [34]. Given that the overwhelming majority of GWAS participants are of European ancestries [34], this lack of portability threatens to exacerbate existing disparities in health outcomes [19] as PGSs begin to see clinical use [34]. Because our framework can jointly model effect sizes across multiple genetically diverse cohorts, it allows information to be shared across cohorts of different ancestries, which we show can increase PGS performance both when applying PGS to ancestry-matched cohorts as well as when porting PGS to cohorts of different ancestries.

Because our framework unifies the inference of the distribution of effect sizes with fitting effect sizes for individual variants, we also examine the inferred distribution of effect sizes for many traits both within a single cohort and across cohorts. We find that commonly-assumed models of effect sizes, such as the point-Normal [59], are badly misspecified; instead, across traits, effect sizes are multi-scale, spanning several orders of magnitude. Standard summaries of effect size distributions, such as the variance of effect sizes or genetic correlation, are sensitive to variants of large effect. Thus, our results call into question the utility of using these measures as adequate summaries of the distribution of effect sizes. We also find that across traits there is no simple relationship between sparsity and the heaviness of the distribution’s tail, highlighting the inadequacy of some recently used models that conflate these two aspects of the distribution of effect sizes with a single parameter [22]. When comparing two cohorts, we find that the inferred effect size distributions have different degrees of correlation at different scales, and appear highly non-Normal, again suggesting that simple summaries of these distributions are inadequate.

Finally, as an application of our framework’s ability to have different priors for different classes of variants, we investigate the relationship between frequency and effect size and find that there is no simple universal relationship between the two. Previous work has assumed that the variance of the effect size distribution for variants of different frequencies, *f*, should scale like [*f* (1 − *f*)]^*α*^ for various *α* typically between − 1 and 0 [28, 46, 64]. In contrast, we find that while rarer variants tend to have larger effects, this general rule does not hold for all traits, but *α* ≈ − 0.4 provides a qualitative fit to many traits.

We have implemented our model in a software package called *Vilma*, which is available at https://github.com/jeffspence/vilma.

## 2 Results

### 2.1 A flexible, unified modeling framework for variant effect sizes

We developed a modeling and inference framework linking the effects of individual variants to a learnable distribution of effect sizes. To begin we considered what properties any such framework should have and then worked to build a model with those properties while still being amenable to efficient inference. We posit that any such model should:

- **Have a flexible, learnable prior on effect sizes:** In general, little is known about the distribution of effect sizes for any given trait. Any modeling framework should learn this distribution from the data, and the distributions it can possibly infer must be sufficiently rich to model traits with varying degrees of polygenicity and effect sizes ranging over several orders of magnitude.
- **Properly account for LD:** The marginal effects estimated by GWAS include the effects of linked variants. This means that the effect of an individual variant is essentially double counted. It is counted once in its own marginal effect estimate, but then appears again in the marginal effect estimated at each of its LD partners. To avoid this double counting LD must be taken into consideration and properly modeled.
- **Require only GWAS summary statistics:** Only summary statistics are typically released from GWAS. Any modeling framework should operate directly on summary statistics to avoid requiring access to individual level data.
- **Easily incorporate prior knowledge:** We often have prior knowledge from additional data about which variants are likely to have large effects. For example, we might expect the effect of a variant to differ *a priori* depending on local chromatin context, expression patterns of nearby genes across tissues, or variant attributes such as frequency, LD score, or being a protein coding variant.
- **Be extensible to multiple cohorts and multiple traits:** As dense phenotyping projects across genetically diverse groups become the norm [21, 37, 56], frameworks should be easily to extend to multiple cohorts or multiple traits.
- **Allow for scalable, accurate inference:** Any modeling framework must remain amenable to efficient approximate inference schemes.

We propose a modeling framework that satisfies these design principles (Figure 1). The key components of our framework are 1) the “regression with summary statistics” model [66], which provides a likelihoood for observing a set of GWAS summary statistics given LD data and the true effects of each variant; and 2) a multi-variate extension of the adaptive shrinkage prior [55], which can flexibly model a broad class of distributions, while remaining amenable to efficient inference schemes. Here we focus on the case of a single trait measured in either one or two cohorts.

**Figure 1:**
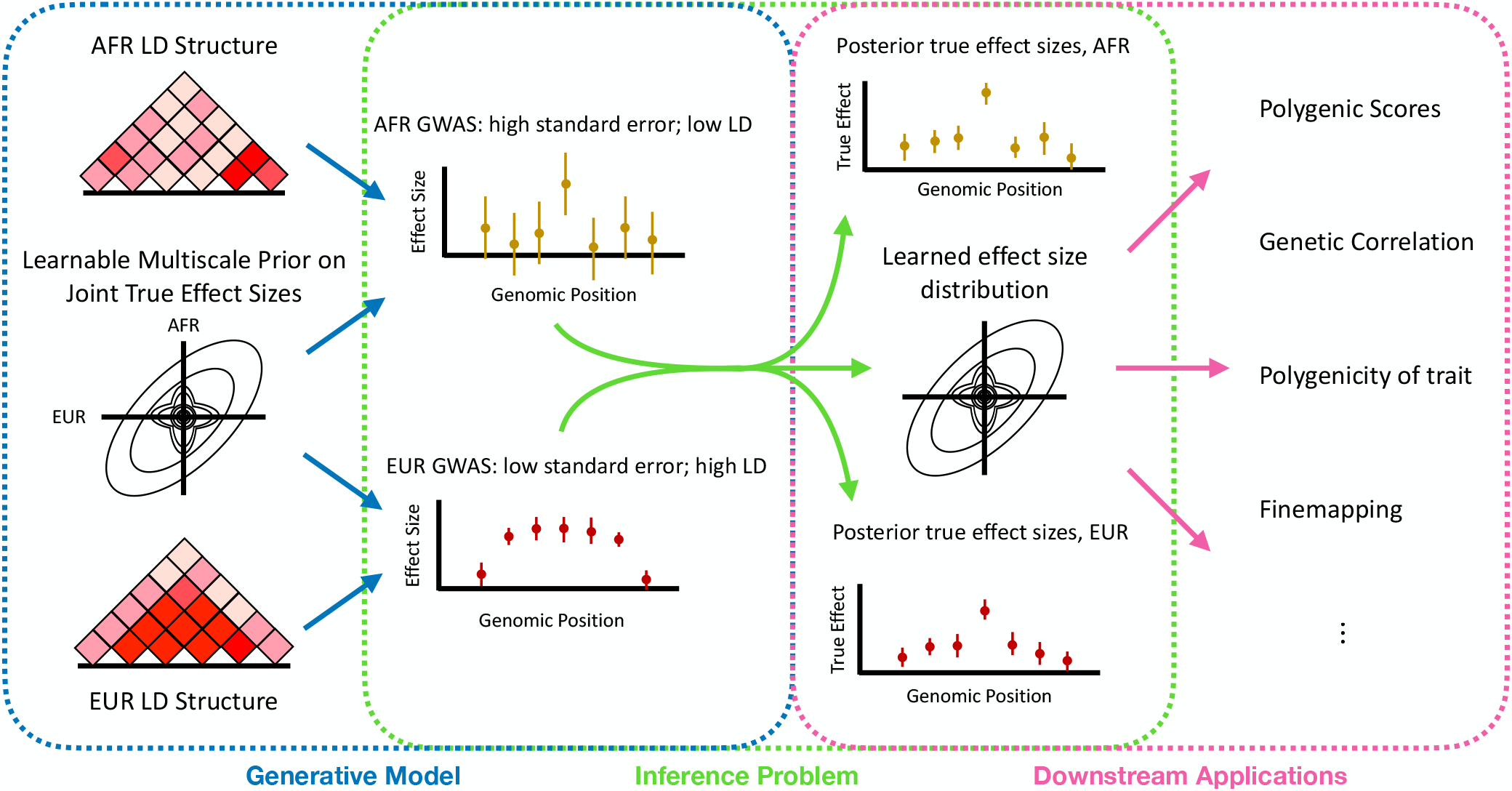
Cartoon of our modeling framework, *Vilma*: *Vilma* models GWAS summary statistics using a learnable prior on effect sizes and estimated LD structure. Using variational empirical Bayes and variational inference, *Vilma* obtains estimates of the distribution of effect sizes as a well as estimates of the true effect sizes. These estimates can then be used for building polygenic scores, measuring genetic correlation or polygenicity, finemapping, or other downstream tasks.

Our framework can model a broad class of effect size distributions by using a dense scale mixture of Normals. That is, we use a mixture distribution with a large number of mixture components, each of which is a Normal distribution centered at the origin but with a different, pre-specified variance. By including a large number of these mixture components and simply varying the mixture weights, we can model a rich class of distributions while essentially only enforcing unimodality and sign symmetry, both of which are biologically plausible. Unimodality with a mode at the origin encodes the intuition that there should be fewer variants of large effect than small effect. Sign symmetry indicates that the distribution of effect sizes is invariant to swapping which allele is labeled as 1 and which allele is labeled as 0, which is sensible as this labeling is somewhat arbitrary. To extend this framework to multiple cohorts, we simply replace the univariate Normal distributions with multivariate Normal distributions, which entails replacing the pre-specified variances with a set of pre-specified covariance matrices. Mathematically, for *P* cohorts, our model takes *K* pre-specified *P ×P* covariance matrices **Σ**_1_, …, **Σ**_*K*_ (we will discuss how we pre-specify these matrices below) and models GWAS summary data as

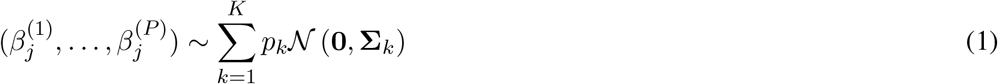

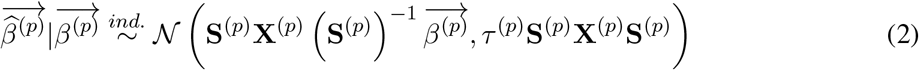

and learns the mixture weights, *p*_1_, …, *p*_*K*_, and GWAS noise “scaling factors”, *τ* ^(*p*)^, for each cohort from the data. We will discuss the inference procedure we employ for this in more detail below. Here, 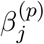 denotes the true effect sizes within cohort *p* at locus 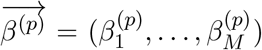 is the vector of true effect sizes across all *M* SNPs in cohort *p*; analogously, 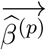 is the vector of marginal GWAS estimates across all SNPs in cohort *p*; **S**^(*p*)^ is a diagonal matrix containing the standard errors of the GWAS estimates in cohort *p* for each SNP along the diagonal; and **X**^(*p*)^ is the LD matrix – a matrix with 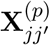 denoting the correlation between the genotypes at SNPs *j* and *j′* in cohort *p*.

Despite the cumbersome notation, the model presented in Equations 1 and 2 has a simple intuitive interpretation. Equation 1 is the joint distribution of effect sizes across cohorts, which acts as a prior on the true (but unknown) effect sizes we see across cohorts at a given SNP. By learning the mixture weights, *p*_1_, …, *p*_*K*_, this distribution is chosen from a rich class of unimodal, sign symmetric distributions to provide an optimal fit to the observed GWAS summary statistics. Equation 2 follows from a central limit theorem type argument on the joint distribution of the marginal effect size estimates and their estimated standard errors (see [66]). Intuitively, the mean of the distribution, 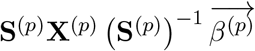 comes from first converting the true effect sizes into *Z*-scores, adding up these *Z*-scores across all SNPs correlated to a focal SNP in proportion to how correlated the genotypes at the two SNPs are, and then converting from this standardized *Z*-score space back to the scale of the original effect sizes to obtain the expected measured marginal effect sizes. Similarly, the variance term comes from noting that the noise in the marginal effect size estimates at two SNPs will be proportional to how noisy the effect size estimates at each SNP are (determined by the frequencies of the alleles at each SNP, which affect the standard errors) as well as how correlated the genotypes are at those two loci (determined by the LD matrix). That is, SNPs with highly correlated genotypes should have highly correlated marginal effect estimates, and unlinked SNPs should have independent marginal effect size estimates. The scaling factor *τ* ^(*p*)^ in our model acts to undo over- or under-correction of population structure, effectively scaling all of the standard errors in a particular GWAS by a constant factor, analogous to the intercept term in LD Score Regression [10].

We can easily extend this model to place different priors on different SNPs (e.g. by allele frequency or functional annotations). Instead of having a single *p*_1_, …, *p*_*K*_ to determine the prior, we simply partition SNPs into several classes and learn a different set of mixture weights per class.

Having formulated our model, we need to be able to efficiently perform inference on it. In particular, we need to fit the hyperparameters *p*_1_, …, *p*_*K*_ and *τ* ^(1)^, …, *τ* ^(*P*)^, and infer a posterior over 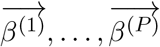. While previous approaches have used cross-validation or a held out validation data to set model hyperparameters [22], these approaches unfortunately require access to individual level genotype and phenotype data, negating the applicability gained by modeling summary statistics as opposed to individual level data. Instead, we would want to take an empirical Bayes approach, where we would set these hyperparameters by maximizing the likelihood of the observed data after marginalizing out the unobserved true effect sizes. Intuitively, this approach treats inferring the distribution of effect sizes from the GWAS summary statistics as its own maximum likelihood estimation problem.

Unfortunately, marginalizing over the unobserved true effect sizes is analytically and computationally intractable. One approach uses the fact that for fixed hyperparameters, Markov chain Monte Carlo (MCMC) can be used to obtain an approximate posterior over the true effects, and given that approximate posterior, it is feasible to maximize a particular function to obtain updated hyperparameter estimates. Alternating these steps of MCMC and updating the hyperparameters is called Monte Carlo Expectation Maximization (MCEM), which approximately finds a local maximum of the likelihood of the data with respect to the hyperparameters. Unfortunately, the need to repeatedly run MCMC makes MCEM notoriously slow. Furthermore, the correlations between genotypes at nearby SNPs either make the MCMC mix slowly or requires costly block updates [22] making it difficult to infer the posterior over the true effect sizes even when the hyperparameters are fixed.

We take an alternative approach, variational inference (VI), that solves both of these problems – setting the hyperparameters via maximum likelihood, and obtaining a posterior over the true effect size. We provide more details in Appendix D, but briefly, VI fits an approximate posterior by minimizing a discrepancy between that approximate posterior and the true, unknown posterior [6]. This turns a computationally difficult sampling problem, MCMC, into the more tractable optimization problem of choosing the parameters of the approximate posterior that minimize this discrepancy. This optimization problem can be solved using standard approaches like coordinate descent or gradient descent. Additionally, the hyperparameters appear in this optimization problem, so we can also minimize this discrepancy between our inferred approximate posterior and the true posterior with respect to our hyperparameters. It turns out that this is equivalent to maximizing a lower bound on the likelihood of the data after marginalizing out the unknown true effect sizes, so this approach is similar to standard empirical Bayes, but instead of maximizing a likelihood, we are maximizing a lower bound on that likelihood [7].

Both MCMC and VI infer approximations of the posterior and in cases where MCMC does not mix well its approximation can be quite poor. VI has been benchmarked in similar contexts [11, 54] where it has been shown to obtain point estimates that are of comparable accuracy to MCMC but using a fraction of the compute budget.

In Section 4.1 and Appendix D we discuss implementation details of both the model and the inference scheme. We also perform a thorough study of the impact of these design choices in Appendix A. Briefly, for the results presented in the main text we consider either one or two cohorts. When there is one cohort, we choose *K* to be 81, and Σ_1_, …, Σ_*K*_ are scalars, so we set them to be approximately uniformly spaced on a log-scale over a data-driven estimate of the likely range of effect sizes. When there are two cohorts we choose *K* to be 144, and since Σ_1_, …, Σ_*K*_ are now matrices, we must specify a variance within each cohort as well as a correlation. We again using a gridding approach, approximately spacing the variances as in the single cohort case, and then uniformly grid across correlations from -0.99 to 0.99. These choices of *K* are somewhat arbitrary and we show in Appendix A.2 that the performance of our method does not depend heavily on the precise number.

We approximate the LD matrix for each cohort by dividing the genome into approximately independent LD blocks [3], and using a low rank approximation to the LD between all SNPs within each block, and setting the LD between SNPs in different blocks to be zero. This approximation results in both computational and memory savings, and has been suggested as a technique to “denoise” LD matrices when using out-of-sample LD [49]. To solve the optimization problem in the VI framework, we perform up to 1000 rounds of coordinate descent, potentially stopping earlier if a round of coordinate descent does not change any of the posterior mean true effect sizes by more than 10^−6^ or if the evidence lower bound (an affine scaling of the objective that we are optimizing that provides a lower bound on the likelihood of the data after marginalizing out the unknown true effect sizes) does not improve by more than 0.1 log-likelihood units.

### 2.2 Application to Polygenic Scores

Polygenic scores (PGS) are a medically relevant use case of the modeling and inference framework presented in the previous section. Under an additive genetic model (evidence for which is discussed extensively in [47]), we could predict the phenotype of individual *i* in cohort *p* with genotypes *G*_*i*1_, …, *G*_*iM*_ across the *M* SNPs as

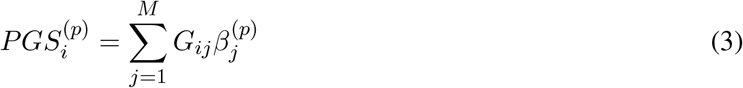

if we knew the true effect, 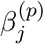, of each variant. Since we do not know these true effects, classical Bayesian decision theory indicates that substituting the posterior mean of the unknown true effects into Equation 3 is the optimal point estimate of Equation 3 in a particular sense [60]:

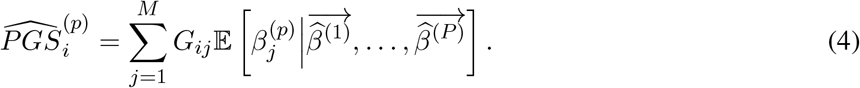

This indicates that by using our modeling framework and extracting the posterior mean effect sizes we can obtain PGSs that are approximately optimal in a specific sense under the assumption that effect sizes are drawn from the distribution that we learned from the data.

In theory we should include as many SNPs as possible in this modeling framework to build the most accurate PGSs. In practice, including additional variants introduces additional computational burden, and some variants have imputation and data quality issues that can result in worse performance. Throughout we use approximately one million variants from the HapMapIII project [15], but explore the impact of the variant set in Appendix A.3.

#### 2.2.1 *Vilma* builds state-of-the-art polygenic scores

To assess the utility of this modeling framework in the context of PGSs we used data from a number of traits from the UK Biobank (UKBB) [56], Biobank Japan (BBJ) [37], and the Million Veteran Program (MVP) [21]. The UKBB is a cohort of individuals living in the UK, with (depending on the exact trait considered and after pruning to unrelated individuals) approximately 320,000 white British individuals, 24,000 individuals of other European ancestries, 6,000 individuals of African ancestries, 7,000 individuals of South Asian ancestries, and 1,000 individuals of East Asian ancestries. The BBJ cohort is a cohort of approximately 140,000 mainly Japanese individuals. MVP is a cohort of veterans of the United States armed forces including approximately 60,000 African-American individuals in the version 2 release used here. We had individual-level access to the UKBB, and so we used the white British individuals for building PGS and tested the accuracy of the PGS in the other sets of individuals. For MVP and BBJ we only used GWAS summary statistics, and for some of the following analyses used those in combination with summary statistics from the GWAS on white British individuals. Across the different cohorts we considered various subsets of 37 blood and urine biomarkers as well as standing height and BMI. Details about cohort delineations, phenotype definitions, and GWAS details are presented in Section 4.2.

We used our modeling framework and Equation 4 to build PGSs using these GWAS data. To begin, we used only the white British individuals from the UKBB, and considered the performance of this PGS in a standard use case: applying the PGS to an “ancestry-matched” cohort, for which we used the individuals of other European ancestries in the UKBB. To assess how well our PGSs perform compared to existing methods, we compared to PRS-CS, which has previously been shown to be the state-of-the-art method for PGS construction using data from a single cohort [22]. Up to some technical details discussed in more detail in Section A.4, PRS-CS uses the same likelihood as our model (Equation 2), but instead of having a learnable prior like our Equation 1, PRS-CS uses a continuous mixture of zero mean univariate Gaussians with the mixture weights coming from a particular fixed distribution that is chosen to induce sparsity. Whereas our model learns the entire unimodal distribution of effect sizes, PRS-CS has a single learnable hyperparameter, which can either be learned from the data (by placing a somewhat informative fixed prior on it and obtaining a posterior) or can be tuned using a validation set of individual level data. Since the main appeal of modeling summary statistics instead of individual level data is to avoid needing access to individual level, we compared our method against the version of PRS-CS that learns its hyperparameter from the data.

Depending on the trait, our framework either performs comparably to PRS-CS or substantially better in terms of Pearson’s correlation *r*, and the squared correlation, *r*^2^, which measures the amount of phenotypic variance explained by the PGS, as a measure of predictive performance. The results are presented in Figure 2a. Across traits this performance is statistically significant (*p* ≪ 10^−16^; two-sided meta analysis over traits; see Section 4.3 for statistical details). While much of this improvement derives from two traits related to bilirubin, the increase in performance remains significant when restricting to the remaining traits (*p* = 3.2 *×* 10^−10^). Bilirubin is an unusual trait in that there are several linked variants each with very large effects. Consistent with this observation, we looked at features of the learned effect size distributions across these traits, and we found that our modeling framework significantly outperforms PRS-CS on traits where *Vilma* predicts that there are several variants with extremely large effects, which we will discuss further below. This highlights the utility of having a flexible, learnable prior – *Vilma* can learn that while there are many variants of small effect, there can still be a few variants with effect sizes that are orders of magnitude larger. In contrast PRS-CS has a single learnable hyperparameter that must simultaneously fit the distribution of effect sizes across multiple scales, necessarily trading off accuracy in fitting one part of the distribution with fitting another part.

**Figure 2:**
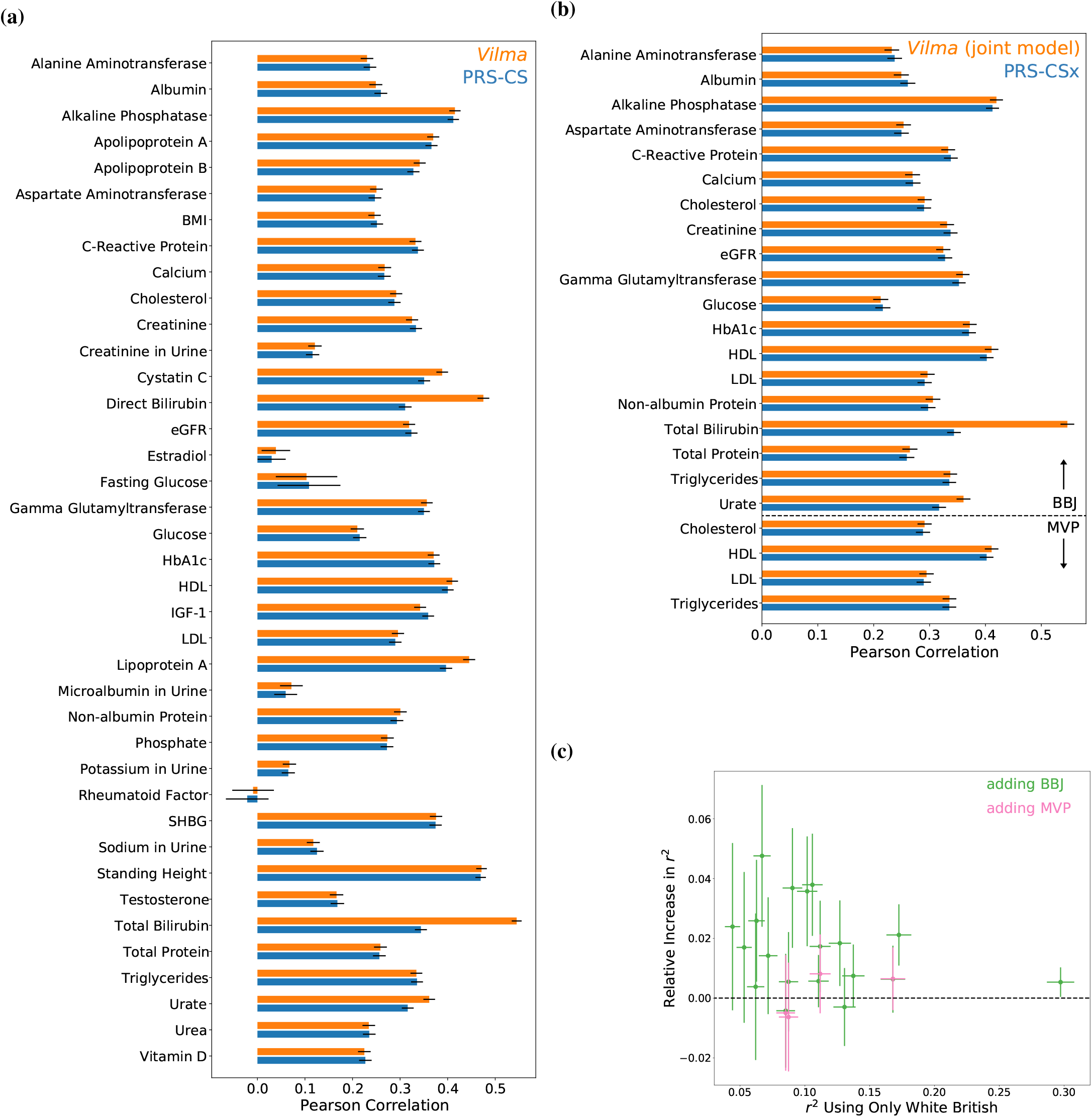
Polygenic score performance in an “ancestry-matched” cohort: **(a)** Pearson correlation between European ancestry individuals’ true trait levels and PGS constructed by either *Vilma* or PRS-CS using white British individuals from the UKBB. **(b)** Same as (a) but using information from both white British individuals from the UKBB and either the BBJ cohort or African Americans from MVP. **(c)** Relative increase in *r*^2^ when comparing *Vilma* PGS built using both white British individuals and either BBJ or MVP individuals to PGS built using only white British individuals.

We next investigated the improvement in prediction accuracy from modeling multiple cohorts. Given that many traits have been estimated to have high genetic correlations across cohorts of different ancestries, we expected that jointly modeling cohorts should improve effect size estimation [9]. We compared our method to PRS-CSx, an extension of PRS-CS that jointly models multiple cohorts, and which is the only other method that performs joint inference of effect sizes across cohorts from summary statistics while properly accounting for LD [45]. PRS-CSx assumes that the magnitude of effect sizes is similar across cohorts, but curiously (and undesirably) assumes that the genetic correlation of the trait across cohorts is zero. For example, given that a variant has a large trait-increasing effect in one cohort PRS-CSx assumes that the effect will also be large in the other cohort, but assumes that it is equally likely that the variant increases or decreases the trait. A model similar to that used in PRS-CSx was also used in the context of inferring whether variants are causal across cohorts or in a cohort-specific manner [48]. There are, of course other approaches to combine data across cohorts such as meta-analysis (improving marginal effect size estimates by averaging across cohorts, but ignoring LD), mega-analysis (pooling individuals from multiple cohorts prior to performing GWAS, ignoring effect size and LD differences across cohorts), or multi-PGS (taking linear combinations of PGS trained in each cohort separately, which does not share information across cohorts when estimating the effect sizes within each cohort) [32]. Yet, given that these methods are performing fundamentally different tasks, we restrict to comparing *Vilma* against PRS-CSx.

We considered two cases of modeling multiple cohorts. We jointly modeled summary statistics from white British individuals from the UKBB with summary statistics from either African American individuals from MVP or primarily Japanese individuals from BBJ. We again considered the standard use case of applying these PGS to an “ancestry-matched” cohort, assessing PGS quality in held-out individuals of European ancestries from the UKBB. The results are presented in Figure 2b. We see that like in the single cohort case, across traits, *Vilma* either performs comparably to PRS-CSx or provides substantial improvement. Across traits this increase in performance is statistically significant (*p* = 3.0 *×*10^−4^ for MVP; *p* ≪ 10^−16^ for BBJ). This shows that our method is state-of-the-art.

We also wanted to explore how much predictive performance is increased by modeling an additional cohort. We compared PGS built using *Vilma* with a single cohort to PGS built using *Vilma* with two cohorts (Figure 2c). We find that improvement is significant when adding BBJ (*p* ≪ 10^−16^), and increased but non-significant when adding MVP likely due to only considering four traits (*p* = 0.69 adding MVP). In both cases the improvements are generally small (adding BBJ: median relative increase in *r*^2^ = 2.6%; max relative increase in *r*^2^ = 6.2%; adding MVP: median relative increase in *r*^2^ = 1.0%; max relative increase in *r*^2^ = 1.7%). Thus, even by adding an “ancestry-mismatched” cohort of a smaller sample size, we can gain a slight, but significant improvement in PGS performance compared to simply using a single “ancestry-matched” cohort.

In Appendix A we show that *Vilma* is robust to our various design decisions. In particular, we show how *Vilma* performs when we tweak various aspects of the model including the choice of dataset used to estimate the LD matrix (Appendix A.1), how many mixture components we use in the prior (Equation 1; Appendix A.2), which variants are included in the PGS (Appendix A.3), and whether we place the prior on effect sizes in their natural scale or whether we first frequency-scale them by 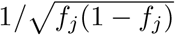 (Appendix A.4).

#### 2.2.2 *Vilma* improves portability

PGSs are known to suffer from poor “portability”: the performance of PGSs significantly degrades when applied to individuals that are genetically not-well represented in the cohort used to build the PGS [34, 5, 42]. Various factors contribute to lack of portability, including differences in allele frequencies, LD, and true effect sizes [62, 40]. Within a cohort, both the accuracy of GWAS estimates and the contribution to predictive accuracy scale with *f*_*j*_(1 − *f*_*j*_) for a given effect size, indicating that variants that contribute the most to PGS predictive accuracy also have the smallest standard errors. In contrast, when we move to a cohort of different ancestries, these become uncoupled, so that the variants that contribute the most to PGS predictive accuracy in the target cohort may not be well-estimated in the GWAS cohort. Similarly, when not all variants are included in the PGS, the effect of a particular variant incorporates the effects of linked variants that are not included in the PGS, and the extent to which these linked effects should be incorporated depends on LD. LD differs between different ancestries so this too can contribute to the portability problem. The true effect sizes may also differ across cohorts due to differences in epistatic or gene-environment interactions [40]. In any case, given that the overwhelming majority of GWAS participants have European ancestries, this lack of portability threatens to exacerbate disparities in standard of care as PGSs see clinical use [34]. Since we saw improved predictive accuracy in an “ancestry-matched” cohort when jointly modeling multiple cohorts, we considered if this joint modeling could also improve predictive accuracy when porting PGSs to a cohort with different ancestries.

We first considered the case of building a PGS in one cohort and then porting it to an “ancestry-mismatched” target cohort. We consider two cases. In both cases we started with a PGS trained using the white British individuals from the UKBB. In one case we compared the performance on individuals with African ancestries in the UKBB to the performance when jointly modeling the UKBB white British individuals with African Americans from MVP. In the other case, we compared the performance on individuals with East Asian ancestries in the UKBB to the performance when jointly modeling the UKBB white British with the BBJ cohort.

Across traits we can see (Figure 3a) the portability problem in PGSs built using just the white British. In European ancestry individuals, *r*^2^ is much higher than it is in other target cohorts (across traits, 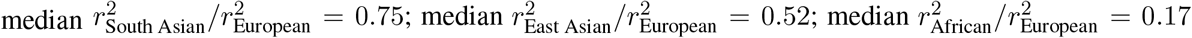) and meta-analyzing across traits the drop is significant for porting to any cohort (*p* ≪ 10^−16^). Yet, we also see (Figure 3b) that including an “ancestry-matched” cohort substantially improves *r*^2^ in the target cohort (median relative increase in *r*^2^ = 11.8% in East Asians, *p* = 5.9 *×*10^−12^; median relative increase in *r*^2^ = 48.0% in Africans, *p* = 1.3 *×*10^−9^). Indeed, the median ratio of *r*^2^ in the target cohort compared to *r*^2^ in European ancestry individuals increases from 0.52 to 0.65 in East Asians and from 0.14 to 0.22 in Africans when including an additional cohort, even after accounting for the improved performance in European ancestry individuals that we saw in Figure 2c.

**Figure 3:**
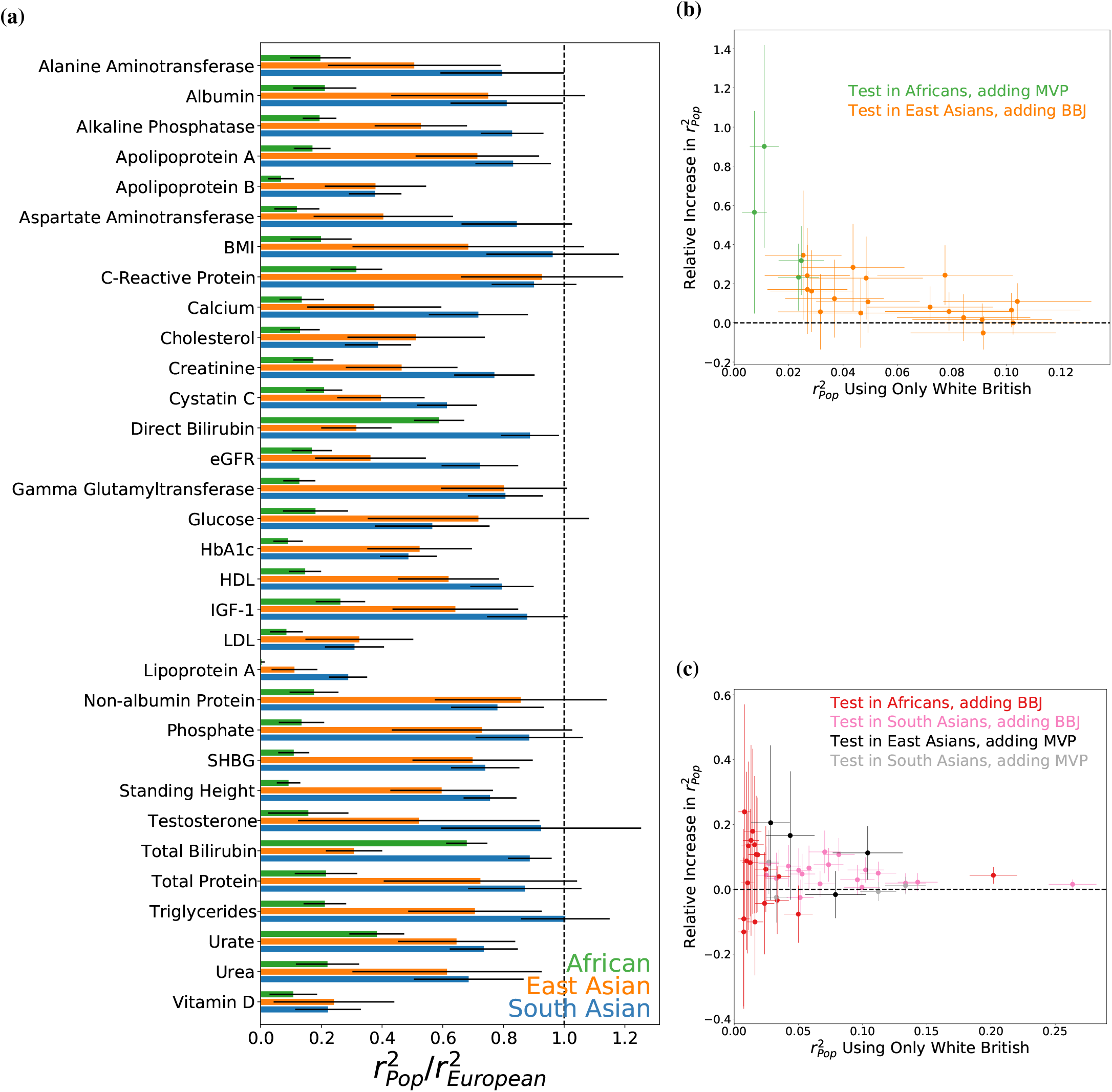
Portability of polygenic scores: **(a)** Squared Pearson correlation, *r*^2^, between true trait values in an “ancestry-mismatched” target cohort and the PGS predicted by *Vilma* using the white British individual from the UKBB relative to the *r*^2^ for individuals of European ancestries. **(b)** The relative increase in *r*^2^ for PGS built using *Vilma* when adding an “ancestry-matched” cohort. That is, for East Asian ancestry individuals we compare a PGS built using white British individuals and the BBJ cohort to one built using just the white British; for African ancestry individuals we perform the same comparison using African Americans from MVP. **(c)** Relative increase in *r*^2^ when adding an “ancestry-mismatched” cohort.

We also looked at the portability problem when neither modeled cohort is “ancestry-matched” to the target cohort. In particular we looked at jointly modeling African Americans in MVP with white British from UKBB and applying the PGS to individuals of East Asian or South Asian ancestries in the UKBB or jointly modeling the BBJ cohort with white British from the UKBB and applying the PGS to individuals of African or South Asian ancestries in the UKBB. The results are shown in Figure 3c. While the gains are more modest compared to adding an “ancestry-matched” cohort, we again see substantial improvement by incorporating additional data. When adding BBJ, the median relative increase in *r*^2^ = 10.5% for Africans (*p* = 3.4 *×* 10^−6^), and 6.5% for South Asians (*p* ≪ 10^−16^). When adding MVP, the median relative increase in *r*^2^ = 15.3% for East Asians (*p* = 1.3 *×* 10^−3^) and 1.2% for South Asians (*p* = 0.03). We also see that the *r*^2^ in the target cohort does again improve relative to the *r*^2^ in European ancestry individuals even after accounting for the improved performance in European ancestry individuals (for BBJ, 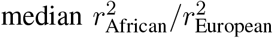 improves slightly from 0.174 to 0.184, and 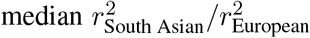 improves slightly from 0.782 to 0.824; for MVP 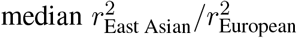 improves from 0.561 to 0.636, but the 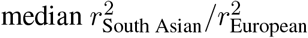 remains essentially unchanged, from 0.586 to 0.585), indicating a slight overall increase in portability across almost all cohorts.

We show in Supplemental Figures A1 and A2 that across all of these portability scenarios we continue to either perform comparably to competing methods or substantially better depending on the trait. Across cohorts and traits *Vilma* performs significantly better than PRS-CS (*p* ≪ 10^−16^), and adding BBJ to *Vilma* outperforms running PRS-CSx with the same cohorts (*p* ≪ 10^−16^), and again *Vilma* outperforms PRS-CSx when adding MVP (*p* = 0.02).

In all cases, we see that jointly modeling multiple cohorts improves predictive performance, whether in an “ancestry-matched” cohort or when porting to a cohort with different ancestries. We also see that modeling multiple cohorts improves the portability of PGSs regardless of whether the additional cohort is “ancestry-matched” to the target cohort.

### 2.3 Estimated Effect Size Distributions

Our framework jointly estimates the effect sizes of individual variants and the overall distribution of variant effects. In the previous section, we considered applying this framework to build PGSs, which relies on the accuracy of the individual variant estimates. Here we turn to the estimated distributions of effect sizes, which are automatically obtained during the course of fitting the models in the previous section.

We can compare these inferred effect size distributions to the distributions that are commonly used as priors in statistical genetics, to test how reasonable such distributions might be. Throughout statistical and population genetics, effect sizes are typically assumed to follow simple distributions such as the Normal, or a Normal with a point mass at zero. An important feature of such distributions is that effect sizes tend to be of a characteristic order of magnitude. That is, most non-zero effect sizes are on the order of one standard deviation away from zero. In contrast, given that different genes may have more or less direct impacts on a trait, and variants can range from slightly perturbing expression to totally disrupting protein function, we might expect from first principles that effect sizes range over many orders of magnitude. Even beyond assuming a particular distributional form, summarizing effect size distributions by their second moments is ubiquitous in statistical genetics. For example, traits are often summarized by their heritabilities when thinking about a single cohort [10] or their genetic correlation across cohorts [9]. Both heritability and genetic correlation are related to the variance (second moment) of the effect size distribution. Given that second moments are dominated by variants of large effect, and effect sizes might span several orders of magnitude, second moments are relatively uninformative summaries multi-scale distributions.

One note of caution in interpreting these inferred effect size distributions is that we only include approximately one million HapMapIII SNPs that pass our filtering criteria [15]. SNPs that are not included in this set but are in linkage with one or more SNPs in this set will have their effect on the trait absorbed into the linked SNPs. Since we account for linkage between the SNPs within our SNP set the effects of these “phantom” SNPs will not be overcounted, but the effects attributed to any single SNP could be an amalgamation of the effect of a variant at that particular SNP as well as some component of linked but untyped SNPs.

We begin by visualizing and summarizing the effect size distributions for various traits in a single cohort. We use the results from models trained using summary statistics from white British individuals from the UKBB. As seen in Figure 4a, the effect size distributions are far from Normal, and across all traits possess some standard features, leading us to posit that these are likely to be universal features of the effect size distribution for sufficiently complex traits. First, the effect size distributions across all traits possess some mass near zero, but a significant amount of mass away from zero, suggesting that at least for this variant set, many – but far from all – variants contribute to each trait. Second, the effect size distributions are multi-scale in that they have substantial mass across multiple order of magnitude, suggesting that there are sets of variants with different “scales” of effect sizes. Finally, these two properties appear to be relatively uncoupled: the percentage of variants with essentially zero effect does not strongly depend on how many variants we would expect under the learned prior to have an effect of at least 0.1 standard deviations of the phenotype (Figure 4b).

**Figure 4:**
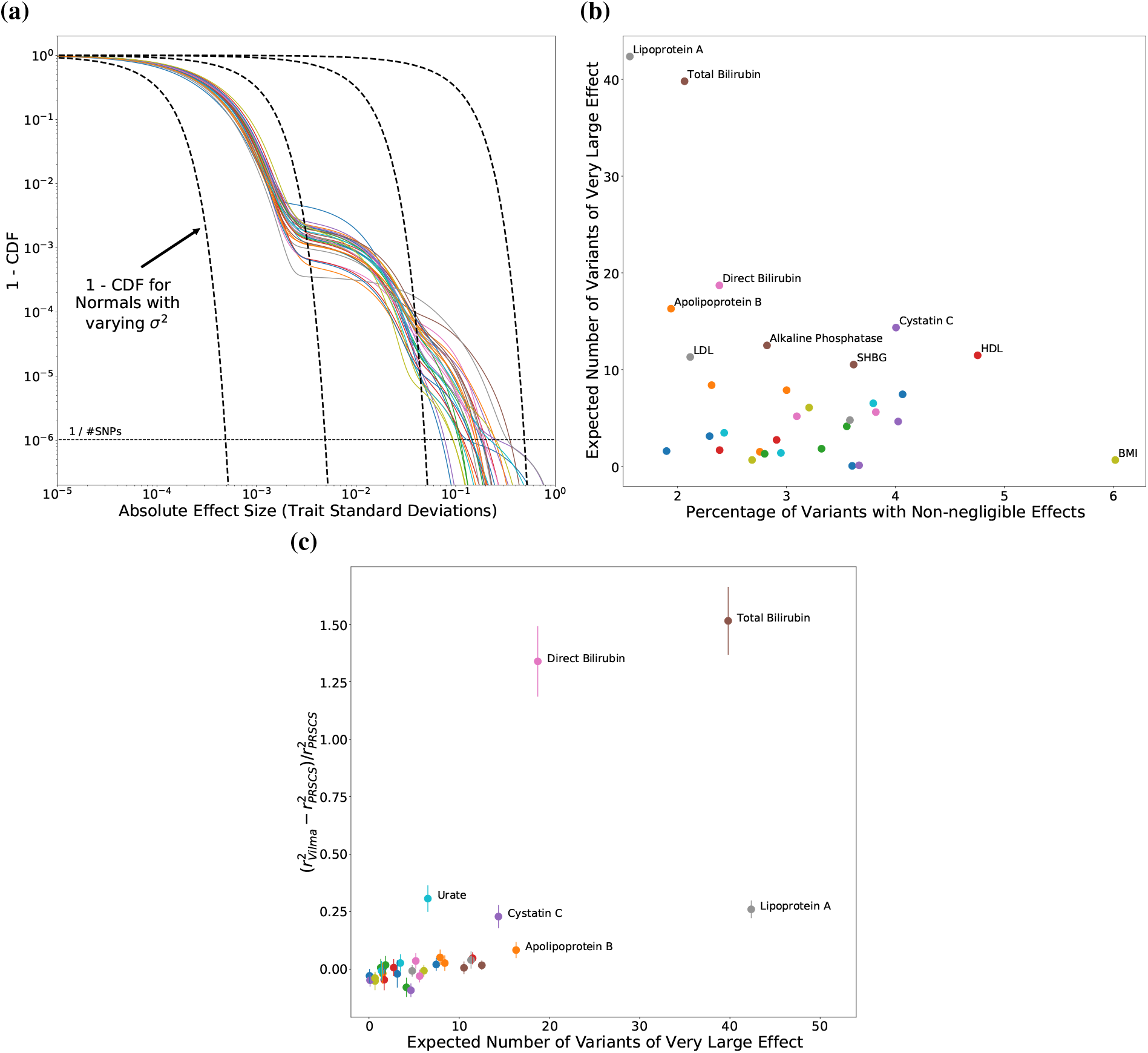
Learned effect size distributions: **(a)** Inferred effect size distributions, plotted as 1 − CDF, which is ℙ (|*β*_*j*_|*> b*) as a function of *b*. All traits where *Vilma* produced a PGS with *r*^2^ *>* 0.02 are plotted. Overlaid in dashed lines are 1 − CDF under the assumption that *β*_*j*_ is Normally distributed. The dotted line is at 1/#SNPs, providing a cutoff below which we do not expect to find any variants in our SNP set. **(b)** A measure of how heavy-tailed the learned trait distributions are plotted against a measure of the sparsity of the trait. We measure heaviness of the tails using the expected number of SNPs of very large effect (#SNPs *×*ℙ (|*β*_*j*_| *>* 0.1), with a very large effect being greater than 0.1 standard deviations. To assess sparsity we use the proportion of variants with an effect size greater than 0.001 standard deviations. **(c)** The relative improvement in *r*^2^ for *Vilma* PGSs relative to PRS-CS PGSs, both trained using white British individuals form the UKBB and tested on European ancestry individuals in the UKBB plotted against the expected number of SNPs of large effect as in (b). *Vilma* has the largest performance gains relative to PRS-CS for traits where *Vilma* infers that there are many variants with large effects.

Given that *Vilma* substantially outperformed PRS-CS on a handful of traits, we sought to further understand this difference in performance. In particular, we wanted to see if any features of the inferred effect size distribution stand out for the traits where *Vilma* outperforms PRS-CS. We find that for these traits, there are several variants of large effect (Figure 4c), and we hypothesize that since PRS-CS has only a single hyper-parameter that is learned from the data, that hyperparameter must trade off accurately modeling the sparsity simultaneously with modeling variants of large effect. For traits for which there are a large number of variants of small effect, PRS-CS is forced to choose a distribution of effect sizes with an appreciable amount of mass near zero, but this then over-shrinks the variants with large effects resulting in poor performance.

Our modeling framework can also infer flexible joint distributions of effect sizes across cohorts. This allows us to go beyond estimating genetic correlations and begin looking more thoroughly at how effect sizes are shared across cohorts. In Appendix B we investigate some of these joint distributions of effect sizes.

Overall, these results highlight the utility of inferring effect size distributions both for learning about complex trait biology, as well as for improving the accuracy of individual effect size estimates.

### 2.4 Frequency-stratified effect size distributions

How the distribution of effect sizes depends on frequency has been a matter of debate [53]. One set of statistical genetics tools is based on the assumption that all variants are expected to contribute equally to heritability, which is equivalent to assuming that variants with frequency *f*, come from a distribution of effect sizes with variance *σ*^2^/(*f* [1 − *f*]) [10, 59]. This relationship between frequency and effect size distribution is an emergent property for variants of large effect in certain models of stabilizing selection [50]. Another set of statistical genetics tools makes the opposite assumption that all variants have the same distribution of effects. Recent work has suggested interpolating between these two by assuming that conditioned on, *f*, effect sizes come from a distribution with variance *σ*^2^ *×* (*f* [1 − *f*])^*α*^ for some *α*. At *α* = − 1 all variants contribute equally to heritability, and at *α* = 0 effect size and frequency are independent [46, 64]. For most traits *α* lies between these extremes suggesting that neither of the standard models adequately describe empirical observations [28, 46, 64].

Our modeling framework can easily estimate different distributions of effect sizes for different sets of variants, and so we investigated the relationship between frequency and effect size distribution by binning variants by frequency. By comparing effect size distributions across these frequency bins, we can begin to more thoroughly explore the relationship between effect size and frequency, as opposed to summarizing this relationship by a single parameter.

For the results presented here, we group variants by their minor allele frequency quintile. For the set of variants we considered this resulted in minor allele frequency breakpoints of ≈ 9%, 18%, 28%, and 39%, and approximately 200 thousand variants per bin. We also considered using 10 or 50 bins. The results were qualitatively similar to those discussed below.

Across traits we find that lower frequency variants do tend to have larger effects, but the story is much more complicated than a single parameter would suggest, as can be seen for three representative traits in Figure 5a. This is especially the case for the variants with the largest effects, where frequency bin is less predictive of overall effect size.

**Figure 5:**
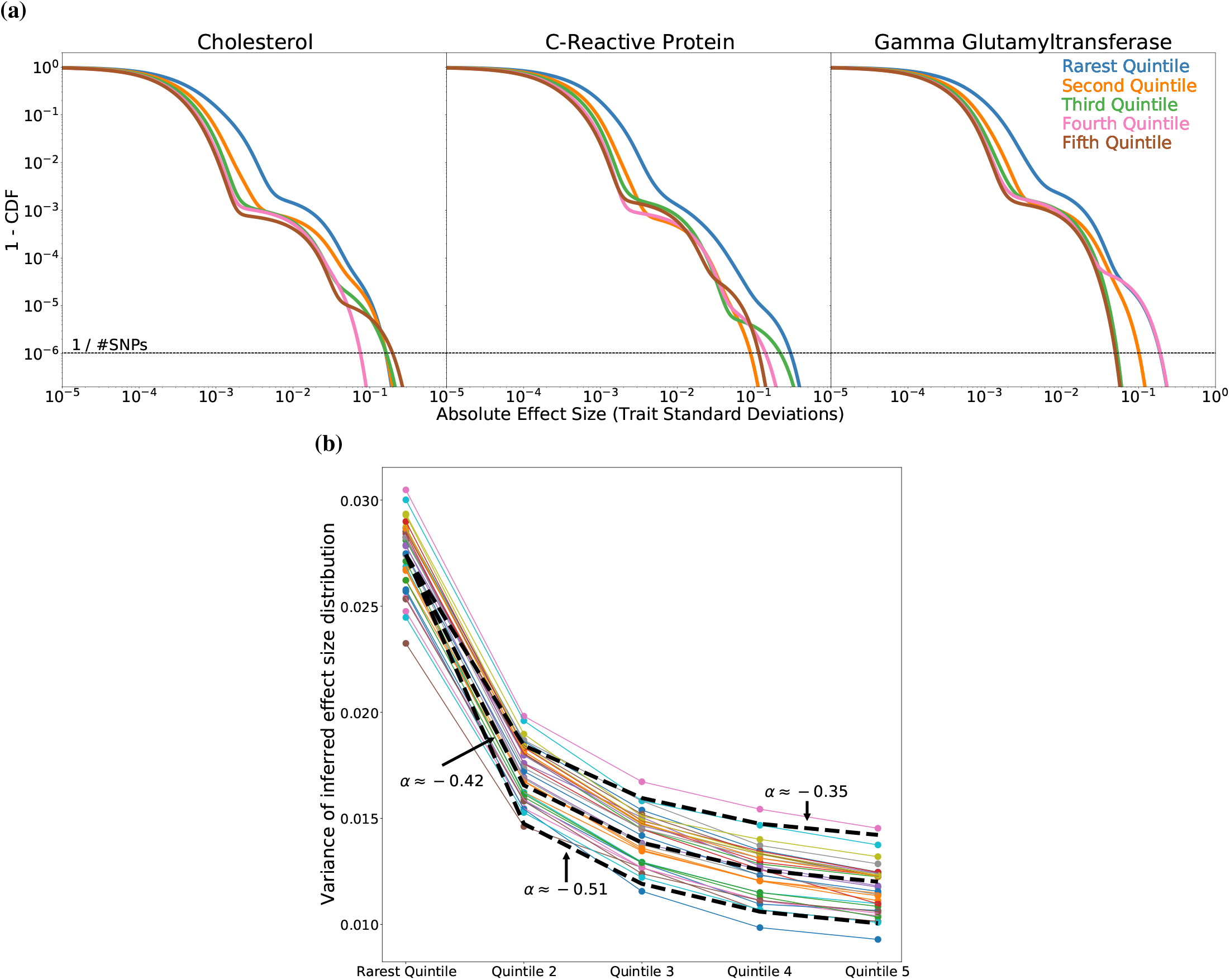
Frequency dependence of effect sizes: **(a)** Effect size distributions for variants in different minor allele frequency quintiles as encoded in 1 − CDF for three representative traits. The dashed line is at 1/#SNPs, providing a cutoff below which we do not expect to find any variants in our SNP set. **(b)** Variance of effect size distributions across minor allele frequency quintiles for all traits where *Vilma* produced a PGS with *r*^2^ *>* 0.02 (colored lines). The black dashed line is the prediction from an *α* model with *α* = − 0.42, which provides a qualitative fit to the behavior of the effect size variance-frequency relationship across traits. We note that variance is not a good summary of the distributions in **(a)** due to their heavy tails.

We also looked at how the variance of the effect size distribution depends on frequency given its prominence in previous models. The results are shown in Figure 5b and show that while the variance consistently increases for rarer variants, this relationship varies by trait to some extent. An *α* model with *α* = − 0.42 is qualitatively similar to the behavior across traits, but does not perfectly describe all traits. This is in line with previous estimates of *α* across a broad range of traits [28, 46, 53, 64]. Note that this only shows a qualitative fit of the *α* model to the *variance* of the effect size distribution as a function of frequency. We reiterate that these heavy-tailed distributions are poorly summarized by their variances.

## 3 Discussion

Here we presented a flexible modeling framework to tie the distribution of effect sizes in one or more cohorts to the summary statistics obtained from GWAS. This framework infers the overall distribution of effect sizes, but also estimates the effects of individual variants while properly accounting for LD. We showed the utility of our approach on two downstream tasks: building accurate, portable PGSs, and investigating the genetic architecture of complex traits.

Our method improves PGS performance. In particular, our results show the importance of flexible priors. Our results also show that including information from multiple cohorts can improve PGS performance whether applying the PGS in an “ancestry-matched” cohort, or when “porting” it to a cohort with different ancestries. This increase in performance is most substantial in cases where the additional cohort used in the model is “ancestry-matched” for the target cohort. Taken together, these results highlight the utility of our model and its ability to incorporate information from multiple cohorts, but also highlight the importance of collecting genotype and phenotype data from diverse cohorts. Finally, in Appendix A.4 and Appendix A.5 it is clear that at least in terms of PGS prediction using common variants, the relationship between effect size and allele frequency is not strong enough to be important for PGS with current datasets. That is, the predictive benefit of flexibly modeling the frequency dependence of the effect size distribution does not outweigh the statistical cost of inferring separate effect size distributions across frequency bins.

In the context of building PGS using data from multiple cohorts, it should be noted that our modeling framework infers effect sizes for each cohort. Here we always used the effect sizes estimated for the white British cohort, but deciding which effect sizes to use in practice to build a PGS for a particular individual is an interesting open problem. Previous approaches, including PRS-CSx, use a validation dataset to learn a linear combination of multiple PGSs [32, 45]. Such approaches could certainly be applied to the output of *Vilma* in a straightforward fashion, so we did not explore them here. These approaches do have a few undesirable features. First, requiring an individual-level validation dataset limits the utility of directly modeling only GWAS summary statistics, and even if individual-level data is available, it is unclear how to optimally split those individuals into a GWAS cohort for estimating effect sizes and building PGSs and a validation cohort for learning hyperparameters and how to weight component PGSs. Second, such approaches assume that when the PGS should be applied to a particular individual, its accuracy on that individual is well-estimated by its performance in the validation cohort. However, genetic ancestry is continuous in nature, whereas PGSs built in different cohorts must necessarily treat those different cohorts as discrete entities. As such, the optimal weighting of component PGSs may vary from individual to individual in a way that is predictable from their genetics. In any case, determining how to optimally combine the multiple PGSs output by *Vilma* or PRS-CSx is an interesting area for future study.

Beyond PGSs, we used our framework to infer effect size distributions and found that the architecture of complex traits is more complicated than previously assumed. Effect sizes are universally multi-scale so standard models such as Normal or Normal plus a point mass at zero are grossly misspecified. Even less standard distributions such as that used in PRS-CS cannot capture the complexity of the distribution of effect sizes with a single hyperparameter. A simple example is that we observe that the sparsity and heaviness of tails of the distribution of effect sizes can independently vary across traits, so no family of distributions determined by a single parameter can simultaneously fit both of these important features of the underlying effect size distribution. Furthermore, the multi-scale nature of effect size distributions calls into question the utility of using summaries based on second moments of the distribution – like heritability or genetic correlation – to compare traits. These measures are sensitive to the behavior of variants of large effect and may not be indicative of the behavior of the vast majority of variants.

We also investigated how the distribution of effect sizes depends on frequency. This relationship is complicated and varies from trait to trait, although in general it does seem that rarer variants tend to have larger effects. Yet, given the multi-scale nature of these distributions, it seems inadequate to summarize this relationship using a single parameter such as *α*. These results highlight the importance of considering variants of large effect when thinking about evolutionary models and the interplay between effect sizes, natural selection, and allele frequencies [16, 28, 50, 63].

Given the simplicity and extensibility of our framework, there are a number of natural avenues for future work. Here we explored inferring different distributions of effect sizes for variants with different frequencies, but it would be trivial to extend this to other functional categories as has been done in different contexts, such as grouping variants by the cell type in which they are active, or whether they lie in enhancer-like or promoter-like regions and so on [18]. Beyond discrete annotations, the distribution of effect sizes for a particular variant could depend on covariates, and this relationship could be modeled via regression, deep learning, or any other machine learning method, and the parameters of such a model could be obtained in a variational empirical Bayes framework similar to how we infer the mixture weights in Equation 1.

We also focused on the case of jointly modeling multiple cohorts, but it would be possible to share information across multiple traits instead of or in addition to multiple cohorts. Some care in the likelihood model needs to be taken in the case where the same individuals were used in each trait’s GWAS. In such a case, the likelihoods of the different traits (analogous to Equation 2) would no longer be independent as an individual’s value of the different traits may be correlated due to correlated measurement or environmental noise. Fortunately, this can be relatively easily fixed as has been done in other contexts [58].

Another potential direction for future work would be to apply *Vilma* to multiple cohorts with similar ancestries, but different environments. Here we considered genetically differentiated cohorts, but recent work has shown that PGSs have poor portability even within an ancestry group when comparing cohorts with different socioeconomic statuses [36]. Similarly, many traits are highly but not perfectly correlated between males and females [4], suggesting that it may be beneficial to consider GWAS separately in males and females and then jointly analyze the results using our framework.

Throughout this work we divided individuals into discrete cohorts as a statistical modeling convenience, but this obscures the fact that no human populations exist in the sense of discrete, non-interacting, panmictic groups [35]. Beyond this, even within a single cohort there will be heterogeneity in individuals’ local environments with consequences for the genotype-phenotype relationship [36]. As noted above, our method can be applied to any set of cohorts regardless of the genetic ancestries or local environments of the individuals involved although we expect there to be more statistical gains when the cohorts are differentiated genetically or environmentally. But we again emphasize that this grouping is a modeling convenience and not indicative of the existence of discrete ancestry groups.

We presented a unifying framework for jointly estimating the genetic architecture of a trait and the effect sizes of individuals. Applying this framework to building PGSs we found that our framework improves over the state-of-the-art method. We also found that across all of the traits we considered the distribution of effect sizes was extremely heavy-tailed, and that the relationship between frequency and effect sizes is much more complicated than commonly assumed in existing methods.

## 4 Methods

### 4.1 Vilma

In this section we describe the model behind *Vilma* and a number of implementation details.

#### 4.1.1 Model

It has been derived previously that the GWAS marginal effects are approximately multivariate Normal conditioned on the true effects [66]:

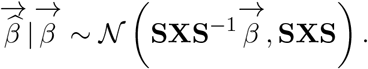

A first thought might be to estimate the true effect sizes using maximum likelihood. After a few lines of algebra, one obtains the maximum likelihood estimator (MLE)

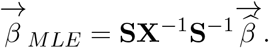

From a frequentist perspective, the variance of this estimator, however, is

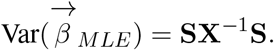

Importantly, SNPs in tight LD cause **X** to have extremely small eigenvalues, which in turn cause **X**^−1^ to have extremely large eigenvalues. This means that the MLE will be extremely noisy, translating into poor predictive performance. Even if we limit ourselves to independent SNPs, we see that the noise is proportional to the squared standard error. This implies that including SNPs for which the standard error is larger than the true effect size will introduce more noise than signal resulting in a worse estimator. Given that much of the signal in many complex traits is explained by SNPs with small effects [8] the MLE is forced to either throw out much of the signal to avoid the attendant noise, or introduce so much noise as to lose any benefit of including those SNPs. To get around this issue, we can place a prior on the true effect sizes, which will regularize our estimates, preventing the variance from exploding.

Unfortunately, it is difficult to model the distribution of true effect sizes. Little is known about how this distribution should look, and for arbitrary complex traits there is no way to use first-principles arguments to derive a simple distribution of effect sizes. A sensible approach would be to try to directly infer the distribution from the data, but then we must decide the class of distributions over which to search. This introduces a key tension in this setup: on one hand, little is known about this effect size distribution and so we would like to make few or no assumptions; on the other hand, if we allow the prior distribution to be arbitrarily flexible there will not be enough regularization resulting in a poorly conditioned and noisy estimator. We therefore must make some assumptions, and we make two simple and sensible assumptions. First, we assume that the distribution of true effect sizes is unimodal, which just means that we expect large effects to be more rare than small effects. Second, we assume that the distribution of true effect sizes is symmetric, which is sensible given that *a priori* we have no reason to suspect that a particular allele will have a particular directional effect. The first assumption always seems sensible, but the second assumption may need to be relaxed in future work if additional information such as local chromatin state or affect on protein coding sequence is incorporated into the model.

Given that we seek to only enforce unimodality and symmetry, we use the adaptive shrinkage prior [55]. The main idea is that many symmetric, unimodal distributions can be approximated by a scale mixture of Gaussians. Concretely, consider this hierarchical construction of a Gaussian scale mixture prior for the true effect size at site *j, β*_*j*_:

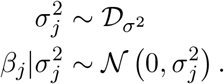

We may approximate a wide class of symmetric, unimodal distributions centered at zero by varying the mixture distribution over the variances, 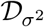, an arbitrary distribution over the positive real numbers.

We may therefore consider an idealized version of our problem as follows:

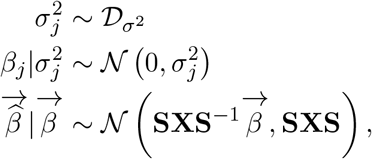

We seek to solve two problems. First, we need to find the distribution 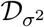 that maximizes the likelihood of the data. Then, with our estimate of 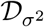 in hand, we can obtain a posterior over *β*. This immediately runs into a practical consideration. If we allow 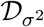 to be arbitrary, then we must infer an arbitrary *function* which will require us to infer and store an infinite number of parameters to represent the function. Instead, following [55], we make a further approximation of discretizing the values that *σ*^2^ can take, considering only a finite number of possible values. We fix this discretization, and then we simply need to infer the mixture weights for this distribution. Concretely, we can consider a set of values of *σ*^2^ – call them 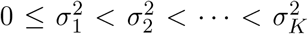 – and we can consider mixture weights Δ = (Δ_1_, …, Δ_*K*_) such that Δ_*k*_ *≥* 0 and 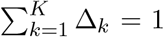. Our model then becomes

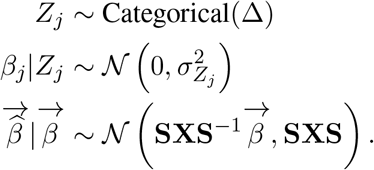

where *Z*_*j*_ acts to index which mixture component a particular SNP draws its effect size from.

Our first problem has now been simplified to finding the *K* dimensional parameter Δ that maximizes the likelihood of 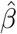.

To extend this model to *P* ≥ 1 cohorts, we note that given the true effect sizes the model of obtaining GWAS data within each cohort remains the same as it only depends on sampling noise that is independent across cohorts. We therefore only need to deal with how to couple the true effect sizes across cohorts. Our generalization is essentially to replace the variances 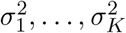 with arbitrary *P × P* covariance matrices Σ_1_, …, Σ_*K*_. Let 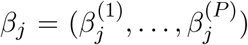 be the true effect sizes in each of the *P* cohorts at position 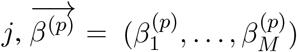 be the *M* true effect sizes across the genome in cohort *p*, and define 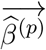 similarly. Let **X**^(*p*)^ be the LD matrix in cohort *p* and **S**^(*p*)^ by the standard errors collected into a diagonal matrix for the GWAS from cohort *p*. Our model is finally:

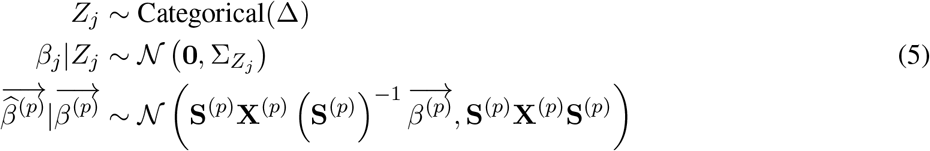

By allowing each Σ_*k*_ to be an arbitrary covariance matrix, our model can capture the genetic covariance between cohorts, which is to say that the model can capture that we expect the true effect sizes to be similar across cohorts. By varying the genetic covariance across Σ_1_, …, Σ_*K*_ we can allow the model to learn the extent to which effect sizes are correlated across cohorts.

We also found a slight but consistent improvement in performance by learning a scaling factor for the standard errors in the model. Concretely, we add an additional hyperparameter per cohort, *τ* = (*τ* ^(1)^, …, *τ* ^(*P*)^) to the likelihood in Equation 5:

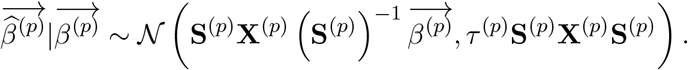

Intuitively, *τ* ^(*p*)^ can account for overcorrection or undercorrection for population structure in the GWAS in cohort *p* analogous to the intercept term in LD Score Regression [10]. We provide additional reasoning behind including these hyperparameters in Appendix C.

With our model in hand, in Appendix D we discuss how to approximately solve the two problems described above: first, optimizing the likelihood over Δ and *τ* to obtain the prior that best fits the data, and second obtaining a posterior over all of the true effect sizes.

#### 4.1.2 Incorporating discrete annotations

Our model can be easily extended to incorporate prior biological knowledge by separating SNPs into different annotations and then inferring annotation-specific effect size distributions. This allows us to formalize the intuition that, for example, SNPs in coding regions might be expected to behave differently on average than SNPs in non-coding regions. To incorporate annotations consider that we have *A* distinct annotations, and a mapping 𝒜 that maps a SNP index to an annotation. That is, 𝒜: {1, …, *M* }→ {1, …, *A* }so that 𝒜 (*j*) is the annotation of SNP *j*. Then, instead of specifying the distribution of effect sizes via a single vector Δ, we have a distinct distribution for each annotation, which we specify via annotation-specific mixture weights Δ^(1)^, …, Δ^(*A*)^. Finally, we replace the prior on *Z*_*j*_ in our model, Equation 5 with

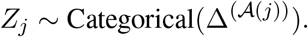

This simple change then results in SNPs with the same annotation having the same prior distribution, with that prior being distinct from the prior on SNPs with other annotations.

In principle one could extend the model to more complex annotations by having a function that maps a SNP index to a vector of mixture weights and then learn that function via empirical Bayes. Such a function could also use additional information about each SNP such as its chromatin state in relevant cell types as a principled way to incorporate such genomics assays into this framework. We leave such an extension for future work.

#### 4.1.3 Computing and approximating the LD matrix

Computing and storing the entire *M ×M* LD matrix **X** would be computationally prohibitive. It would also make the parameter update steps derived in Appendix D take *O*(*M* ^2^) time which would again be prohibitive for the hundreds of thousands or millions of SNPs we considered here. We follow previous approaches [22, 59] by approximating **X** as a block diagonal matrix. In particular, we divide the genome into approximately independent blocks [3] and assume that the LD between SNPs in different blocks is zero. We further approximate the LD matrix by assuming that each each block in the block matrix is low rank. This approximation has been shown to “denoise” the estimated LD matrix when using out-of-sample LD [49], although theoretical justification of this procedure is left for future work. In order to decide on the rank of each block in the block matrix, we perform the singular value decomposition (SVD) and keep only those components with singular values greater than 0.106. This value guarantee that pairs of SNPs with *r*^2^ smaller than 0.8 are guaranteed to be linearly independent in the low rank approximation. In practice, however, many pairs of SNPs with LD much higher than these values can still be linearly independent depending on their values of *r* with other SNPs.

The white British LD matrix–used for GWAS summary statistics derived from the white British cohort of the UKBB–was computed using 10,000 randomly sampled unrelated white British individuals from the UKBB, and the block sizes were determined as in [3] using the 1000 Genomes EUR superpopulation [14]. The European ancestry LD matrix was constructed similarly, but using 10,000 randomly sampled unrelated individuals of European ancestry in the UKBB to compute the pairwise correlations between sites. The African LD matrix–used for the MVP results–was constructed using 6,497 unrelated African ancestry individuals in the UKBB and using block sizes determined from the 1000 Genomes AFR superpopulation. The East Asian LD matrix–used for the BBJ results–was computed using 1,154 unrelated East Asian ancestry individuals in the UKBB and using block sizes determined from the 1000 Genomes EAS superpopulation.

#### 4.1.4 Implementation details and runtimes

Our framework is implemented in a software package, *Vilma*, available at https://github.com/jeffspence/vilma. Inference is heavily optimized using numpy [23] and numba [29], allowing crucial routines to be compiled. We also provide tools for reading and writing PLINK [12, 43] format files containing GWAS summary statistics and constructing LD matrices. For a single cohort and approximately one million variants, *Vilma* runs in a matter of hours using 20 cores. For two cohorts and approximately one million variants, *Vilma* runs in *≈* 30 hours.

### 4.2 GWAS, cohort definitions, and summary statistic acquisition

Genome-wide association studies for serum biomarkers were performed in individual ancestries from the UKBB as previously described [52]. Briefly, individuals in the UKBB were separated by global PCs into European-ancestry (self-identified White British versus self-identified other European ancestries), South Asian ancestry, East Asian ancestry, and African ancestry. Only unrelated individuals were included in the analyses to avoid confounding with family structure. Across all ancestries, biomarker measurements were log-transformed and adjusted for age indicators, sex, fasting time indicators, global principal components, month of assessment, day of sample analysis, and estimated dilution factor of samples. Then, for each ancestry, GWAS were run with genotyping array and within-ancestry PCs as covariates. The For Biobank Japan GWAS, summary statistics were downloaded from JENGER as previously described [26].

For standing height and BMI, individuals with values five or more standard deviations from the mean were removed. These traits were then residualized on age, sex, genotyping array, and the first 18 PCs. We then dropped any individuals whose residualized trait values were five or more standard deviations from the mean and re-residualized, repeating this process until the values converged.

### 4.3 Statistical comparison of PGS

Throughout we compare the performance of PGS by computing the Pearson correlation, *r*, in a set of heldout test individuals. Even for a fixed PGS, the finite sample size of the held-out test set means that for a different test set from the same ancestries we would expect the *r* we calculate in this alternate test set to be somewhat different. In this sense, the *r* that we calculate is an uncertain estimate of some true “population” *r* which would be the correlation across all individuals in the broader population. As such, any difference in performance between two PGSs might be small enough to be due entirely to chance, and which PGS is better may switch on a different test set from the same population. To calculate the statistical significance of the difference in *r* between two PGSs say 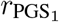 and 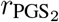 we bootstrap over individuals in the held-out test set and compute 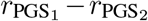 on that bootstrapped dataset. Across bootstraps, this gives us an estimate of the variance of 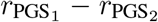, which we can then use to compute a *Z* score for the difference in *r*, assuming asymptotic Normality. When considering multiple traits, we meta-analyze across traits, summing our estimates of the difference in 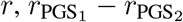 across traits. To obtain the variance of this test statistic we sum the variances of the difference in *r* across traits, which implicitly assumes that the traits are independent. We then convert this statistic to a *Z* score by dividing by the square root of the estimated variance, and compute *p*-values using a two-sided *Z*-test.

We also consider relative improvement in *r*^2^, where we perform the same routine as above, but instead use 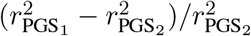 as a test statistic, and use bootstraps to estimate its variance.

To compare *r* across cohorts say 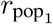 and 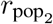 we no longer have to worry about dependencies between the two cohorts, so we compute the variances of 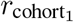 and 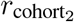 separately by indpenedently bootstrapping the two cohorts. To compute the variance of 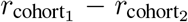 we then add the variances of the 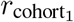 and 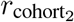 as they are independent.

## Acknowledgements

We would like to thank Fabio Morgante, Matthew Stephens, and members of the Pritchard Lab – particularly Matthew Aguirre, Hakhamanesh Mostafavi, Shaila Musharoff, Roshni Patel, Yuval Simons, and Clemens Weiß – for helpful discussions and feedback. This research was conducted using data from UK Biobank, a major biomedical database (Project #30418 and # 24983). J.P.S. was supported by NIH training grant 5T32HG000044-23. This work was supported by NIH grants R01HG011432 and U01HG012069.

## Appendix A Robustness of *Vilma* and additional results

### A.1 *Vilma* is robust to LD misspecification

By requiring only summary statistics (as opposed to individual-level data), *Vilma* necessarily must make use of LD information. A key assumption underlying the derivation of the likelihood we use is that the LD between SNPs in the sample used in the GWAS is (asymptotically) the same as that in the cohort used to compute the LD matrix [66]. If both cohorts are drawn uniformly at random from the same larger population then this assumption is met. In practice, there are numerous biases in the enrollment of any GWAS cohort which might differentiate it from the cohort used to compute the LD matrix. Furthermore, there may be subtle or substantial ancestry differences between the two cohorts.

To test the extent to which violations of this assumption affect the predictive accuracy of *Vilma* PGSs we compared the results when using three increasingly misspecified LD panels on summary stats derived from a GWAS of white British individuals in the UKBB. First, we used an “in-sample” LD panel constructed using a subset of 10,000 white British individuals. Second, we considered an “ancestry-matched” but “out-of-sample” LD panel, constructed using 10,000 individuals of European ancestries that were not included in the white British GWAS cohort. Finally, we built an “ancestry-mismatched” LD panel, using 6,497 individuals of African ancestries in the UKBB.

When comparing the performance of the polygenic scores constructed using these three LD panels we see virtually no difference between the “in-sample” and “out-of-sample” LD panels. The “out-of-sample” LD panel in fact performs slightly better, but the difference is small (mean increase in *r* across traits and cohorts: 0.001; median increase in *r* across traits and cohorts: 0.0007; *p* = 0.003). There is, however, a substantial drop in performance when moving from an “ancestry-matched” to “ancestry-mismatched” panel (mean decrease in *r* across traits and cohorts: 0.05; median decrease in *r* across traits and cohorts: 0.05; *p* ≪10^−16^). Taken together, this suggests that it is certainly not necessary to obtain an in-sample LD estimate to obtain good performance, but some care should be taken to ensure that the cohort used to estimate the LD panel and the GWAS cohort are as genetically similar as possible. These results are summarized in Figure A3.

### A.2 Performance does not strongly depend on the number of mixture components

The only part of the *Vilma* model that is not fit from the data is the pre-specified grid of covariance matrices. In principle adding additional covariance matrices could slightly improve performance at the cost of additional computational expense – several parts of the *Vilma* method scale linearly in *K*, the number of mixture components. To see if this additional computational cost is worth the benefit we considered performing a gridding of the prior variances, 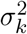, keeping the lowest and highest points of the grid fixed, but changing, *K*, the number of points in the grid (approximately gridding uniformly in log space from the low end to the high end). We considered *K* ∈ {25, 81, 289, 625}, and we found that while there was a slight improvement in performance in going from *K* = 25 to *K* = 81 (*p* = 0.001 in Europeans, but *p* = 0.927 across cohorts), there was no significant improvement in going form *K* = 81 to *K* = 289 (*p >* 0.05 in each cohort, and across all cohorts), and only a tiny improvement when porting to individuals of African ancestries at *K* = 625 (mean increase in *r* of 0.0008 in Africans, *p* = 2 *×* 10^−6^; *p >* 0.05 in each other cohort and across all cohorts). As a result we used *K* = 81 for all of the other single cohort analyses. These results are summarized in Figure A4.

**Figure A1:**
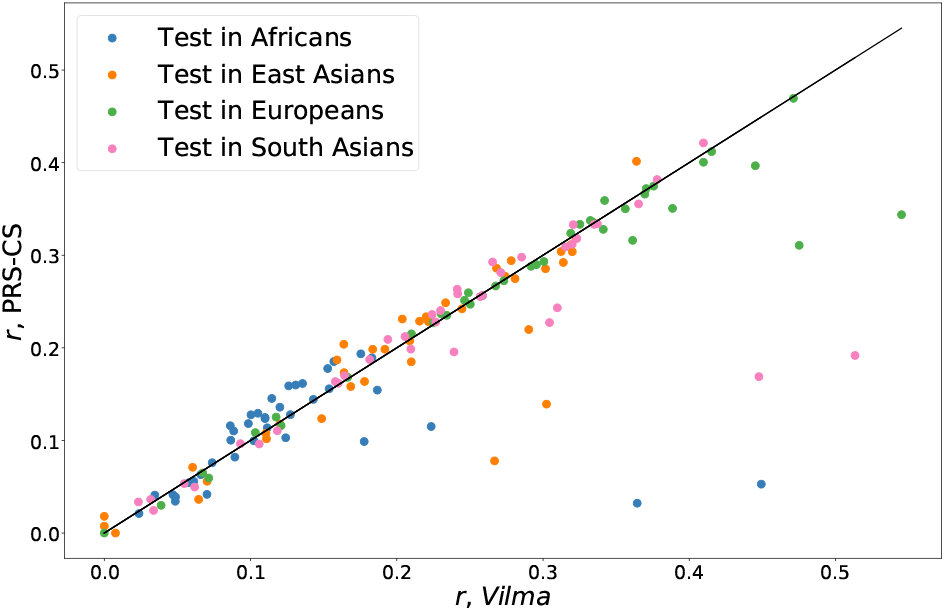
Comparison of *Vilma* and PRS-CS across traits and cohorts. PGSs were built using a GWAS performed on the UKBB white British, and then tested in one of four target cohorts. Each point is a trait in a particular target cohorts of held out individuals in the UKBB.

**Figure A2:**
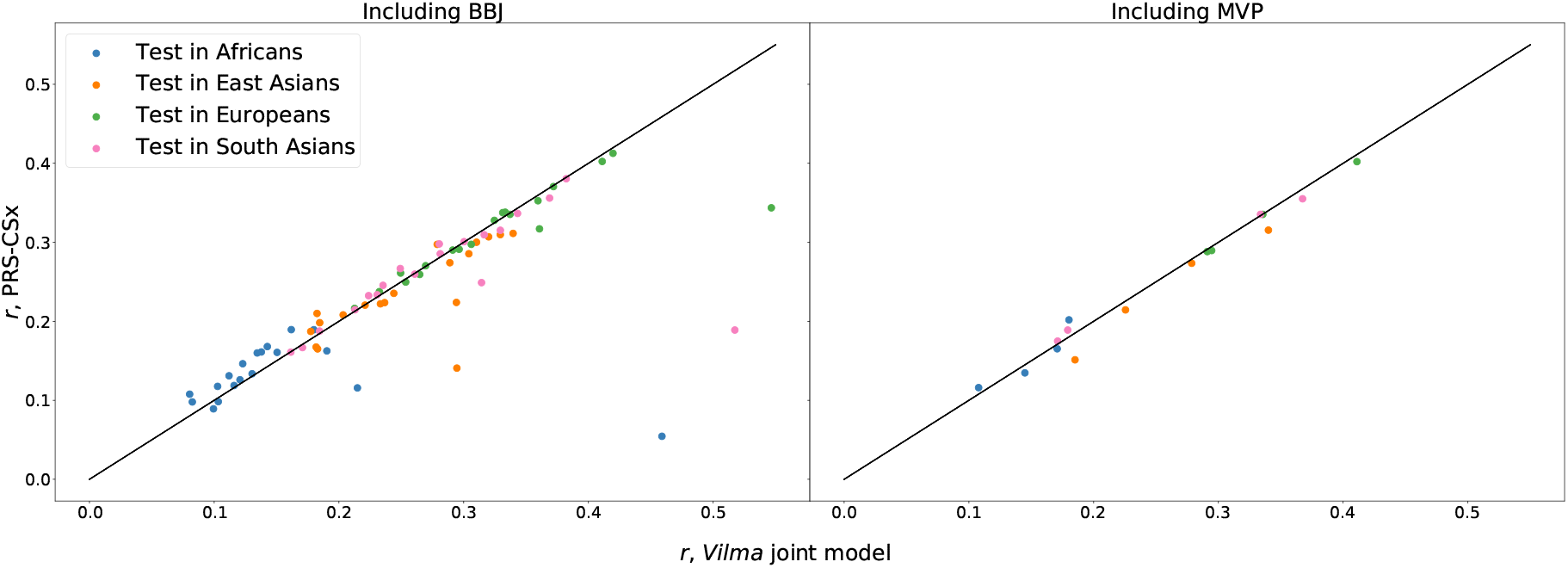
Comparison of *Vilma* and PRS-CSx across traits and cohorts. PGSs were built using a GWAS performed on the UKBB white British combined with a GWAS performed in BBJ (left) or MVP (right), and then tested in one of four target cohorts of held out individuals in the UKBB.

**Figure A3:**
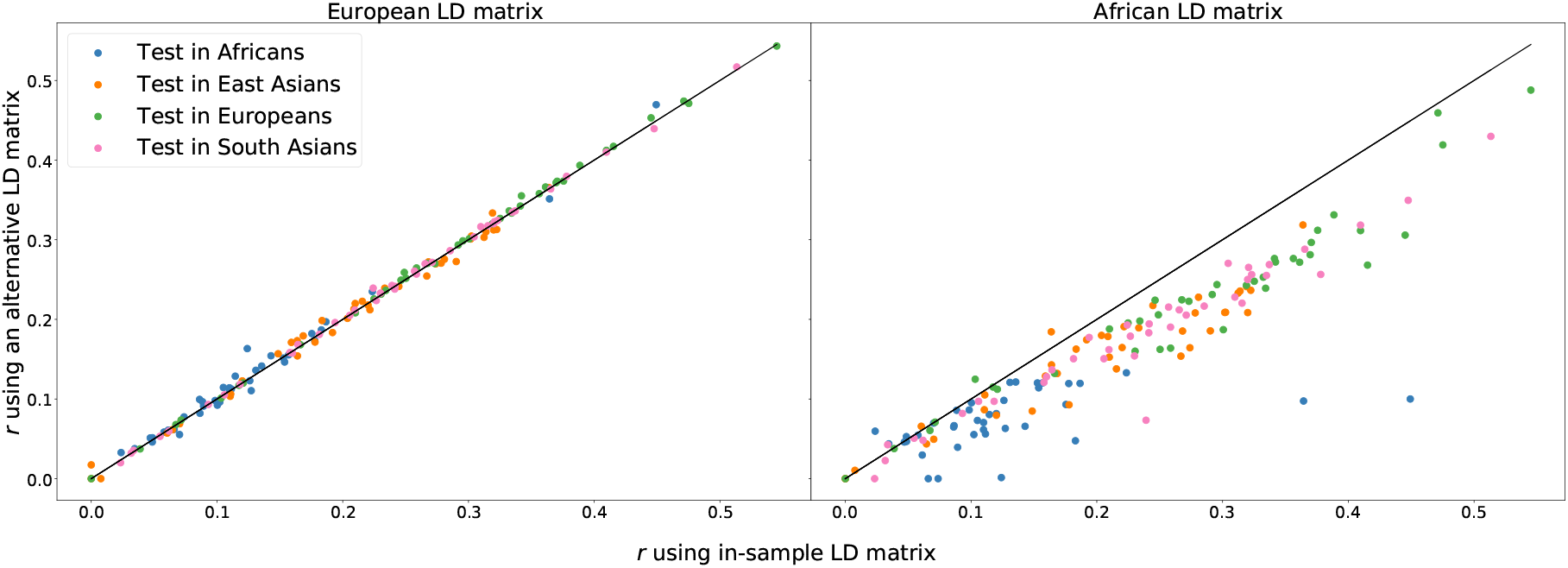
Comparison of *Vilma* PGS performance using different LD panels. The horizontal axis of each plot shows the performance of a PGS built using *Vilma* with an in-sample LD matrix. The vertical axis shows the performance when using either an out-of-sample but “ancestry-matched” sample constructed using held-out individuals of European ancestries (left) or an out-of-sample and “ancestry-mismatched” sample constructed using held-out individuals of African ancestries (right). Each point represents a single trait in a particular held-out target cohort.

**Figure A4:**
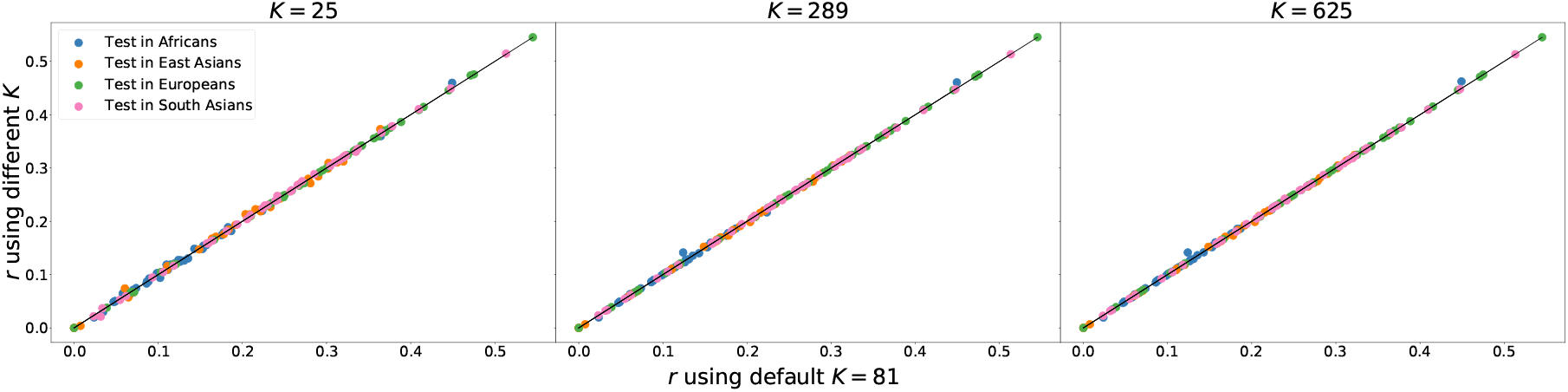
Comparison of *Vilma* PGS performance using different numbers of mixture components. The horizontal axis of each plot shows the performance of a PGS built using *Vilma* with *K* = 81 mixture components – the number used throughout the main text for all single cohort analyses. The vertical axis shows the performance when using a different number of components, either *K* = 25 (left), *K* = 289 (center), or *K* = 625 (right). Each point represents a single trait in a particular held-out target cohort.

### A.3 Using poorly imputed variants can degrade performance

In the main text we exclusively used HapMapIII [15] SNPs that passed our filtering criteria (minor allele frequency *>* 0.001, INFO score *>* 0.3) resulting in a set of approximately one million SNPs. In contrast, we can impute approximately twelve million SNPs with minor allele frequency *>* 0.001 and INFO score *>* 0.3. INFO score is a measure of imputation accuracy and roughly corresponds to the effective proportion of the sample size at that SNP when performing GWAS. That is, for SNPs with an INFO score of 0.5, the power to detect an association is roughly equal to the power in a sample of half the size where that SNP was directly genotyped.

It has previously been reported that variant sets can affect PGS accuracy to some extent for certain methods [42] and we wanted to see the effect on *Vilma*. To explore this, we divided the twelve million SNPs passing our filters into 4 roughly equal quartiles based on their INFO scores. That is, the first quartile contains the approximately 3 million worst imputed SNPs that pass our filters. When comparing the results of *Vilma* using any of these quartile of INFO scores to the results when using the HapMapIII SNPs, we find that there are traits and target cohorts for which one or the other SNP sets perform significantly better than the other. That is, there are always situations in which one quartile outperforms the HapMapIII SNPs and vice-versa. Yet, there are dramatic overall trends as seen in Figure A5. In general, even though the quartiles contain approximately 3 times as many SNPs as the HapMapIII SNP set, when those SNPs are poorly imputed the performance of *Vilma* suffers substantially, which has also been ovserved in [41]. Indeed for the lowest and second lowest quartiles, we see huge drops in performance when looking across traits and target cohorts (*p* ≪10^−16^ in both cases with median drops in *r* of 0.08 and 0.02 respectively). For best imputed and second best imputed quartiles, we obtain slightly better performance than using the HapMapIII SNPs, although it depends on the trait (*p* ≪ 10^−16^ and *p* = 0.00 and median increases in *r* of less than 0.005 in both cases).

### A.4 Model of effect sizes in scaled vs. unscaled space

There is a subtle difference between our model, Equation 5, and several existing PGS models [59, 22, 30]: we place our prior on the effect of each additional allele, whereas other models place a prior on the effect of each addition allele scaled by the standard error of the GWAS estimate of the marginal effect size of that allele. This introduces a dependency between the expected magnitude of the effect size and the frequency of the allele, namely that the variance of the prior for SNP *j* is proportional to [*f*_*j*_(1 − *f*_*j*_)]^−1^. This scaling of effect sizes makes some sense in the single cohort case as this dependence between frequency and effect size is exactly what is expected under a model of stabilizing selection on the trait and a high degree of pleiotropy [50], and it simplifies the math to some extent. Unfortunately, this makes less conceptual sense once we move to more than one cohort – if an allele has different frequencies in two cohorts but the distribution of effect sizes in the scaled space is perfectly correlated across cohorts, then the allele actually has different effects in the cohorts. In a sense the prior assumes that alleles somehow “know” their frequencies in the two cohorts and adjust their effect sizes accordingly. As such we thought it more sensible to place the prior on unscaled effect sizes so that if the distribution of effect sizes is highly correlated across cohorts then alleles actually have similar effects in the two cohorts. Similar considerations arise in the context of genetic correlations. For example see [9].

In any case, we implemented a version of *Vilma* that places its prior on scaled effect sizes and tested it in the single cohort case. The results are presented in Figure A6. Overall we find that while PGS accuracy differs depending on whether the prior is place on scaled or unscaled effect sizes, the differences tend to be small with the unscaled prior performing very slightly better (median increase in *r* of 6.1*×*10^−4^ across traits and target cohorts; *p* = 0.003), and one approach is not clearly better than the other as the best-performing PGS varies from trait to trait. In some sense this is consistent with the results presented in Figure 5 and the following section (Section A.5). Putting a prior on the scaled genotypes is equivalent to assuming that the variance of the effect size distribution scales like [*f*_*j*_(1− *f*_*j*_)]^−1^ whereas putting a prior on the unscaled genotypes is equivalent to assuming that the variance of the effect size distribution scales like [*f*_*j*_(1− *f*_*j*_)]^0^. In contrast, we find that the variance of the effect size distribution scales more like [*f*_*j*_(1− *f*_*j*_)]^−0.42^, highlighting that both priors may be somewhat inappropriate but in different directions.

**Figure A5:**
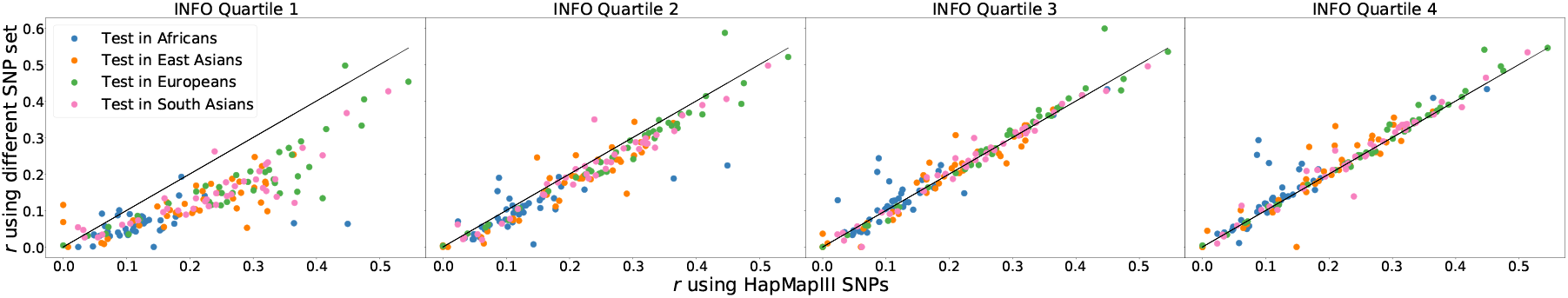
Comparison of *Vilma* PGS performance using different SNP sets. The horizontal axis of each plot shows the performance of a PGS built using *Vilma* with the SNP set used throughout the paper – the SNPs in HapMapIII that have minor allele frequency *>* 0.001 and INFO score *>* 0.3. The vertical axis shows the performance when using a different SNP set. From left to right, the plots show the performance when using quartiles of increasing INFO scores of all approximately 12 million SNPs with minor allele frequency *>* 0.001 and INFO score *>* 0.3. That is, the leftmost plot uses the approximately 3 million SNPs with the lowest INFO scores that pass our filters, the next plot uses the next approximately 3 million SNPs in terms of INFO scores. Therefore the plots are arranged in terms of increasing imputation accuracy. Each point represents a single trait in a particular held-out target cohort.

### A.5 PGS performance when binning by allele frequency

In the main text we discussed models where SNPs in different allele frequency bins had different effect size distributions. In such models we found that rarer SNPs tended to have larger effects, which suggests that PGS performance might improve if we allow different priors in different allele frequency bins. Yet, there is a trade-off here in that fitting additional prior distributions results in less information sharing across SNPs (SNPs with different annotations do not mutually share information) but if the SNPs within an annotation are similar to each other but distinct from other SNPs then the added flexibility may improve PGS performance. As such, we explored this empirically by constructing PGSs in models where we divide SNPs into *B* bins based on their allele frequencies. We considered *B* ∈ { 5, 10, 50}. The results are presented in Figure A7.

Overall, we found that using a single prior across all SNPs very slightly outperformed any model with different priors for different frequency bins (median increase in *r* of 0.008, 0.0015, 0.0018 across traits and target cohorts when comparing a single frequency bin to 5 bins, 10 bins, or 50 bins respectively, with *p* = 0.0009, 0.0003, 0.0002). This indicates that while effect size distributions might change across allele frequencies they do not change by enough to outweigh the additional noise introduced by adding more parameters to the model. This is consistent with Figure 5 in that while there is some signal for differences in distributions across frequency bins, the difference is small, and it seems more important for PGS accuracy to correctly model that multi-scale nature of the effect size distribution than it is to model the relationship between effect size and frequency.

### A.6 PGS performance when keeping *τ* fixed

In Section C we provide theoretical motivation for including a standard error scaling factor *τ* in Equation 2. To see if it actually produces better PGS in practice we compared our standard model to an implementation of our model that keeps *τ* fixed at 1, which assumes that the standard errors are properly scaled. Across target cohorts and across traits, we find that learning *τ* from the data slightly but consistently and significantly improves PGS performance (median increase in *r* of 0.002 across cohorts and traits, *p* = 1.8 *×*10^−6^. The results are shown in Figure A8.

**Figure A6:**
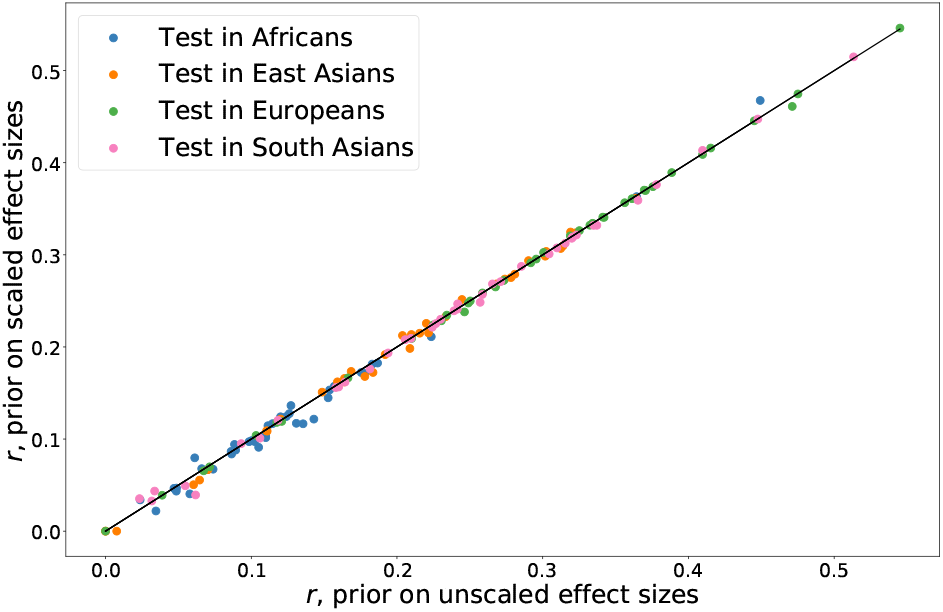
Comparison of *Vilma* PGS performance with a prior on scaled or unscaled effect sizes. The horizontal axis shows the performance of a PGS built using *Vilma* with the default of having a prior on the unscaled effect sizes. The vertical axis shows the performance when instead the prior is placed on frequency-scaled effect sizes as done in many other PGS methods. Each point represents a single trait in a particular held-out target cohort.

**Figure A7:**
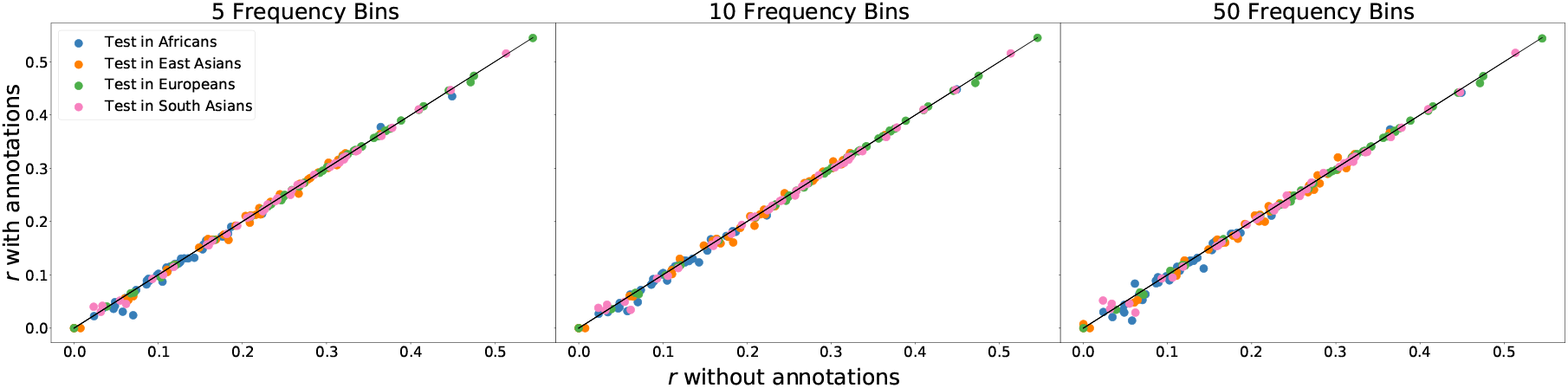
Comparison of *Vilma* PGS performance with varians annotated by various numbers of frequency bins. The horizontal axis shows the performance of a PGS built using *Vilma* with the default of having all variants have the same prior. The vertical axis shows the performance when instead separate priors are learned for SNPs in different frequency bins. We considered either 5 bins (left), 10 bins (center), or 50 bins (right). Each point represents a single trait in a particular held-out target cohort.

## Appendix B Cross-cohort effect size distributions

Our modeling framework can infer flexible joint distributions of effect sizes across cohorts. This allows us to go beyond estimating genetic correlations and begin looking more thoroughly at how effect sizes are shared across cohorts. As such we trained two-cohort models using white British individuals from UKBB and then either African Americans from MVP or the BBJ cohort.

As we see in Figures A9a and A9b the effect sizes are far from Normal with different degrees of correlation emerging at different scales. Comparing representative joint distributions for the UKBB white British and BBJ (Figure A9a) to the joint distributions for the UKBB white British with MVP African Americans (Figure A9b), we see generally higher degrees of correlation in effect sizes between UKBB white British and BBJ than between UKBB white British and MVP African Americans. We infer these effect size distributions using a subset of all SNPs, however, and so effects such as different LD patterns in different cohorts likely play a role in this observation. Finally, we note that there is generally a greater degree of correlation at variants of large effect, suggesting that large, direct effects are more likely to be shared across cohorts than small effects, which may be mediated by more complex pathways allowing for a greater degree of epistatic or gene-environment interactions to result in different effects in different cohorts.

## Appendix C Motivation for the standard error scale factor, *τ*

In the derivation of the likelihood of our model, Equation 2, we implicitly assumed that the squared standard errors from the GWAS can safely be used as plug-in estimates for the true marginal variances. We will show below that this holds approximately for uncorrected GWAS in unstructured populations, but that uncorrected or overcorrected population structure can result in significant deviations between the GWAS squared standard errors and the true marginal variances.

To begin, we consider the usual additive model for the value of the phenotype, *Y*_*i*_, of individual *i* as a function of their genotypes *G*^(*i*)^ = (*G*_*i*,1_, …, *G*_*i,M*_), effect sizes *β* = (*β*_1_, …, *β*_*M*_), and some residual noise *ε*_*i*_. We assume *ε*_*i*_ is uncorrelated across individuals, has mean 0, and variance 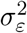.

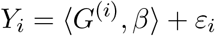

An uncorrected GWAS estimates the marginal effect at SNP *j* as

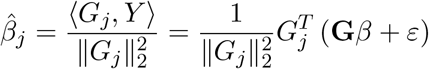

where *G*_*j*_ = (*G*_1,*j*_, …, *G*_*N,j*_) is the collection of genotypes across individuals at locus *j*, **G** is the genotype matrix, and *Y* and *ε* are the trait values and noise terms collected across individuals.

The squared estimator, 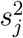, of the standard error of 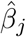 is in turn

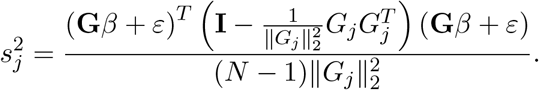

As is usual in asymptotic arguments, we rely on the large sample size of GWAS to assume that 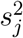 is close to its expected value. We will show that if there is no population structure, then the expected values of 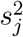 is approximately the true variance of 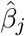, but if there is population structure, then the expected value of 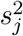 is approximately equal to the true variance of 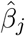 times a multiplicative factor that does not depend on *j*.

**Figure A8:**
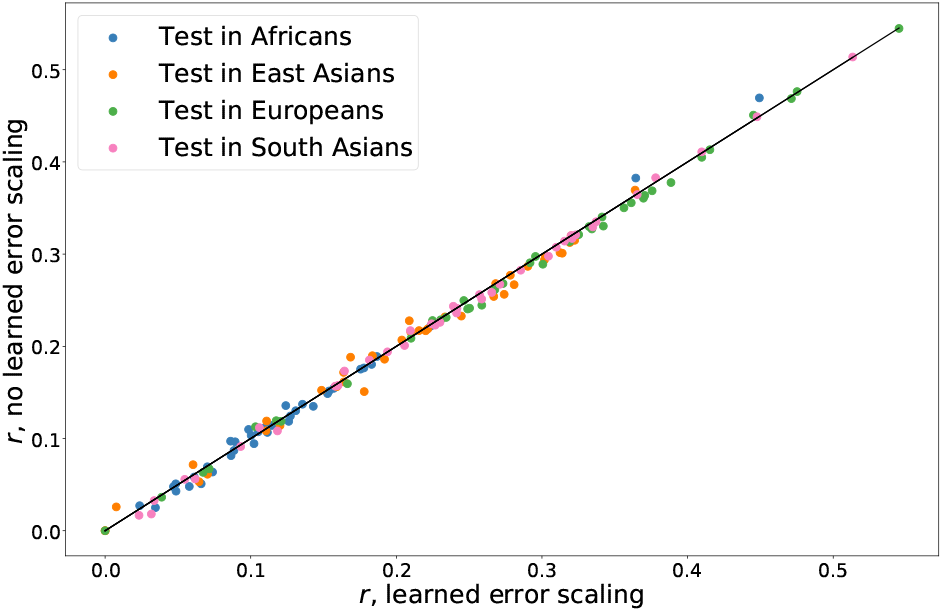
Comparison of *Vilma* PGS performance when learning *τ* or fixing *τ* = 1. The horizontal axis shows the performance of a PGS built using *Vilma* with the default of learning the standard error scale factor, *τ*, in Equation 2. The vertical axis shows the performance *τ* is instead fixed to be 1. Each point represents a single trait in a particular held-out target cohort.

To make this rigorous, we consider the model under which to take expectations. In Section 4.1.3 we discussed how we approximate the LD matrix as being block diagonal, which indicates that under the likelihood in Equation 2, each block is independent. In reality, even SNPs that are in linkage equilibrium will have an in-sample *r*^2^ of approximately 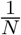 indicating that even though we treat separate blocks as being independent, there is weak correlation between SNPs in separate blocks. While an *r*^2^ of 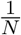 may seem negligible, even in the largest modern studies *M* ≫ *N* indicating that while each individual unlinked SNP asserts a negligible influence on a focal SNP, the large number of unlinked SNPs exert a macroscopic effect on the correlation observed at the focal SNP. Writing *β*^(*b*)^, for the true effects of the SNPs within the *b*^th^ independent block, and assuming that *j* is in that block, we consider

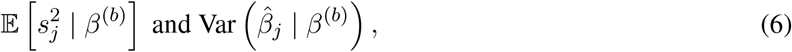

treating the effects of SNPs in different LD blocks as being random effects. In particular, we assume that 𝔼 [*β*_*k*_] = 0 and 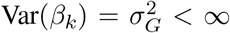 for all *k*, and we assume that *β* and *ϵ* are uncorrelated, but do not make any particular distributional assumptions. To compute the quantities in Equation 6, we will use the notation **G**^(*b*)^ for the genotypes in the *b*^th^ block and *β*^(−*b*)^ and **G**^(−*b*)^ for the true effect sizes and genotypes across the genome exclude block *b*.

**Figure A9:**
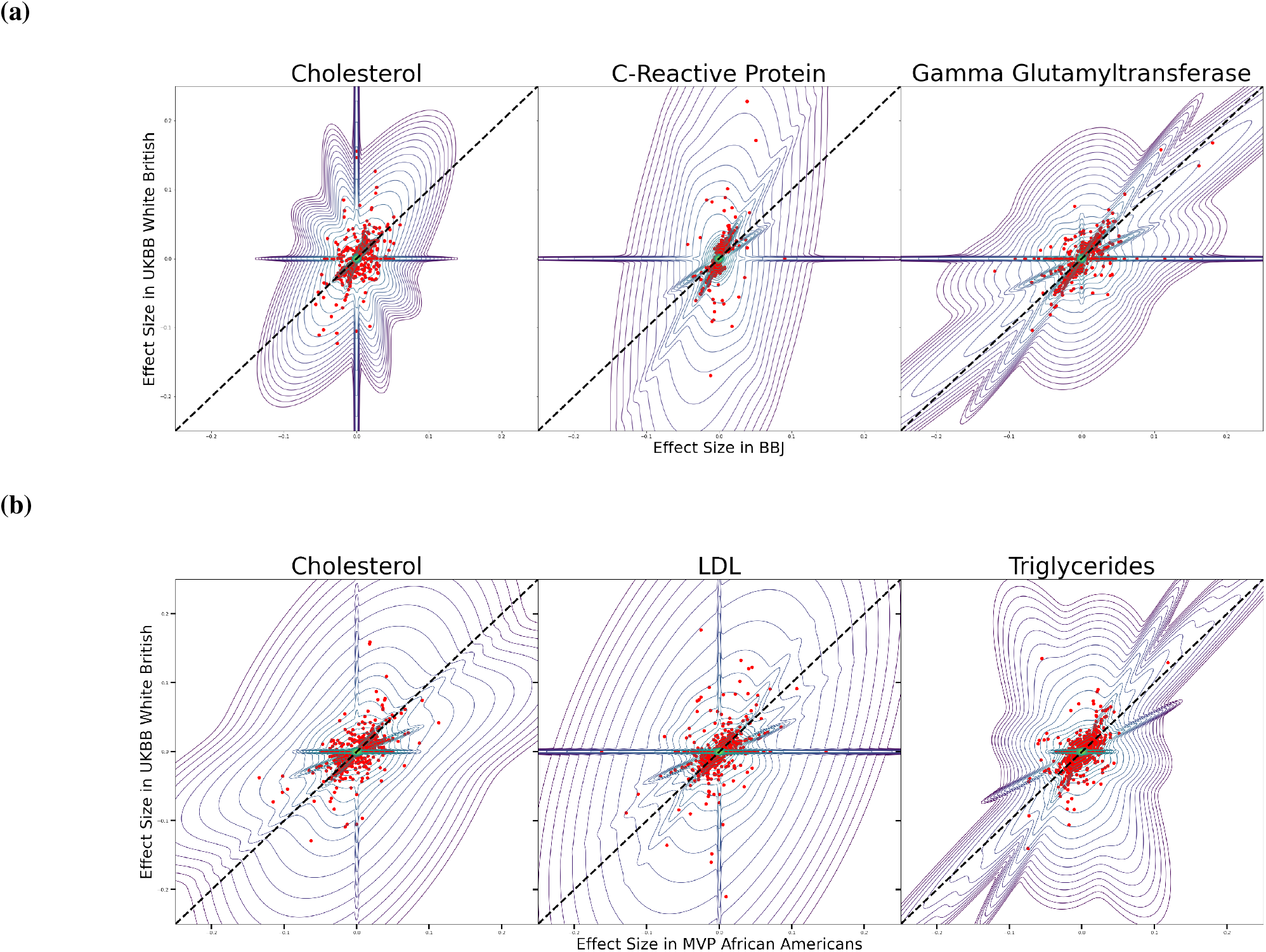
Effect size distributions across cohorts: Inferred joint effect size distributions represented as contour plots learned for three representative traits using data from **(a)** white British individuals in the UKBB and the BBJ cohort or **(b)** white British individuals in the UKBB and African American individuals from MVP. Plots are on a semi-log scale, with effects smaller in magnitude than 10^−2^ being plotted in linear scale and larger effects being plotted on a log-scale.

To begin, we can note that

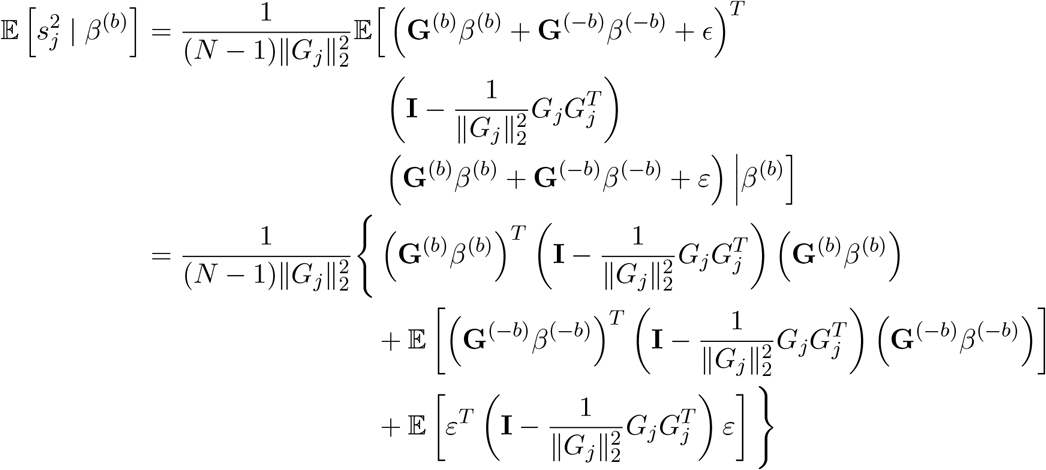

since *β* and *ϵ* are uncorrelated and have mean zero. We assume that block *b* only contains a small fraction of the SNPs and that the true effect sizes in block *b* are not too much larger than what we might expect to see in other blocks. Together these assumptions mean that the term

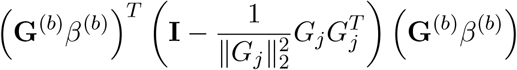

is negligible compared to the other terms. We can now use the formula for expectations of quadratic forms to obtain

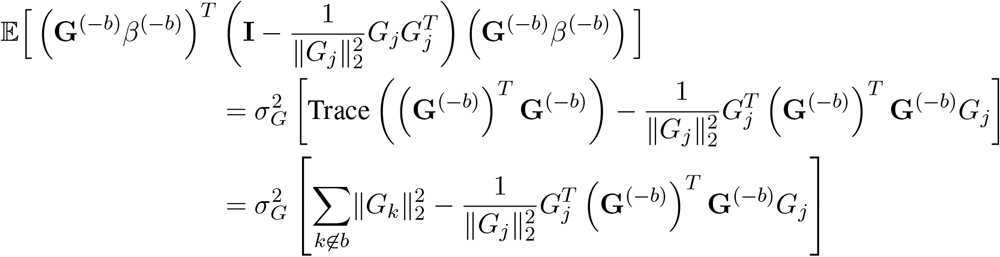

Now, by assumption, block *b* does not contain too many SNPs, and so

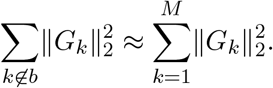

Furthermore, we assume that the *r*^2^ between SNPs in different blocks is approximately 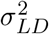. In unstructured populations, 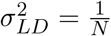, but if structure is overcorrected or undercorrected, then 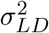 could differ from 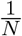. With this assumption,

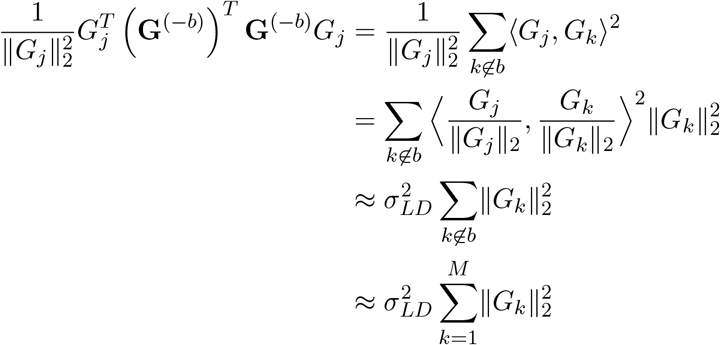

Now, we use that *ε* has mean zero and the formula for expectations of quadratic forms again to obtain

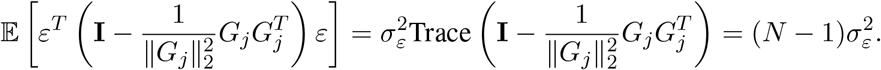

Combining we see

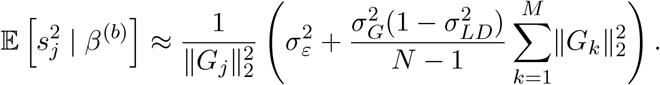

Meanwhile we can readily compute the true variance of 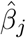 :

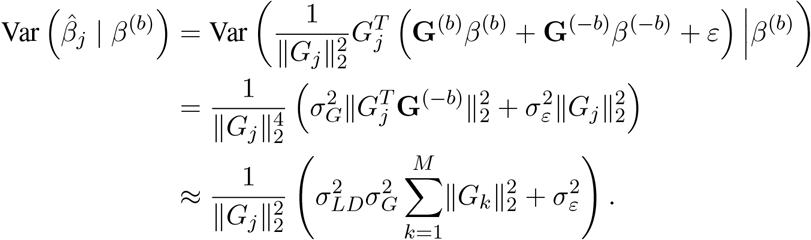

Taking ratios, we see

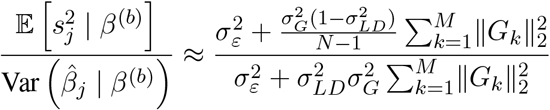

Now, recall that for unstructured populations 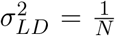, in which case the right hand side reduces to exactly 1 implying that the standard errors are good estimators of the actual sampling variation. On the other hand, if we overcorrect or undercorrect for population structure, then 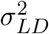 will differ from 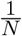 (and possibly be *O*(1)) and the ratio of the expectation of our estimate of the variance to the true variance will not be 1. Importantly, however, this ratio is approximately equal to something independent of *j* suggesting that the estimated variances are off from the true variances by some constant universal factor. We denote this factor in cohort *p* by 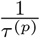. In general we do not know 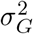 *a priori* and 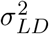 cannot be estimated without access to the genotype data, and so we learn *τ* ^(*p*)^ from the data by treating it as a hyperparameter.

## Appendix D Variational Inference Scheme

Unfortunately our model, Equation 5, is analytically intractable. Throughout this section, we will suppress the vector notation and simply write *β* for 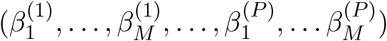 and similarly for 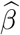. In order to compute polygenic risk scores, we need to be able to compute the posterior mean of 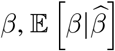. Trouble arises because the *β*_*j*_ are not independent under the posterior. Even in a single cohort, consider two SNPs in strong LD: if the GWAS shows that they are both associated with the trait, it could be that just the first of these SNPs is associated with the trait and the other is only associated through its linkage with the first or *vice versa*. In particular, if it was known that one of the SNPs had a true effect on the trait that explained the GWAS signal at both SNPs, then we would expect the other SNP to not have much of an effect. This non-independence means that the posterior mean for any *β*_*j*_ depends on what is happening at all linked sites. In order to compute the posterior mean we would need to integrate over the value of *Z*_*j′*_ for each of these linked *j*′, which would require *O*(*K*^# Linked SNPs^) time.

Since it is infeasible to obtain the posterior analytically, we must turn to methods to compute an approximate posterior. Classically, posteriors in intractable models are approximated by using MCMC. Yet, MCMC can have trouble mixing, resulting in poor approximations to the posterior, and it can be difficult to assess whether it has converged or not. In the last few decades, VI has become an attractive alternative to MCMC. VI finds an approximate posterior by optimizing an objective function that is equivalent to minimizing the Kullback-Leibler divergence (KL)

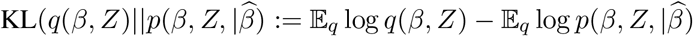

between the density of a variational posterior, *q*(*β, Z*) and the density of the true posterior, 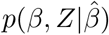 [6]. Minimizing this KL divergence turns out to be equivalent to maximizing a lower bound on *p*(*β*), the **e**vidence **l**ower **bo**und (ELBo):

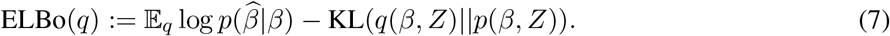

This optimization problem, in turn, can be more tractable, leading to fast algorithms for fitting complex models. There are concerns that VI finds lower quality posteriors than MCMC, but both methods find approximate posteriors, and the posterior mean under the variational posterior is often very close to the true posterior mean even if other aspects of distributions differ. VI schemes for models similar to those described here have been shown to approximate the posterior mean at least as well as MCMC [11, 54], indicating that our use of VI is justified.

The key to speeding up variational inference is defining the family of distributions over which to search for the approximate posterior. By requiring the variational posterior to respect certain independence assumptions, we can avoid the exponential runtime of computing the true posterior. This improved computational performance comes with a statistical price, however: additional independence assumptions can only degrade the quality of the variational posterior. A common approach is to make the “mean field” assumption that all variables are completely independent under the variational posterior [6]. In our case, we can still derive efficient updates making a slightly less draconian independence assumption. We only make the mean field assumption across SNPs – this allows us to capture the dependency in the posterior across cohorts at a SNP. Concretely, we assume that the posterior factorizes as

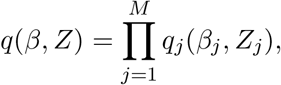

which assumes independence across SNPs, but not across cohorts.

We then assume that each of these *q*_*j*_ is an indexed mixture of Gaussians:

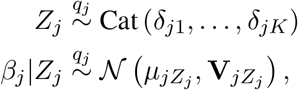

so that for the *k*^th^ mixture component, *β*_*j*_ is Normally distributed with mean *µ*_*jk*_ and covariance matrix **V**_*jk*_. In this formulation *δ*_*j*1_, …, *δ*_*jK*_ are the mixture weights for the *K* different Gaussian components.

One way to think of this is as a mixture of *K* distributions, where for each component distribution *β*_*j*_ is a Gaussian and *Z*_*j*_ is fixed to take a particular, distinct value between 1 and *K*. If we take the mixture weights to be *δ*_*j*1_, …, *δ*_*jK*_, then this exactly matches the above. Furthermore, each of these components is an exponential family and they are non-overlapping in the joint space of *Z*_*j*_ and *β*_*j*_. One way to see this is that *Z*_*j*_ is fixed to be a distinct value in each component of the mixture, so one component cannot put mass on the same part of the joint space of *Z*_*j*_ and *β*_*j*_ as another. Hence, the results of [54] show that this indexed mixture of Gaussians forms an exponential family with natural parameters

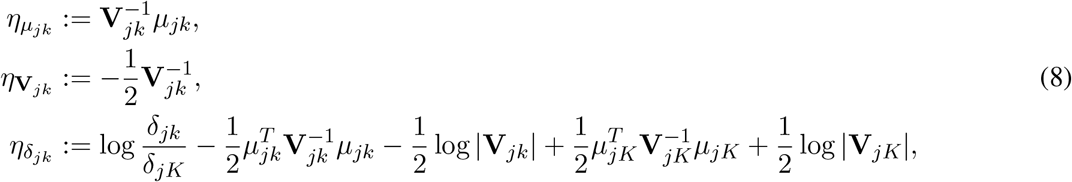

and corresponding sufficient statistics:

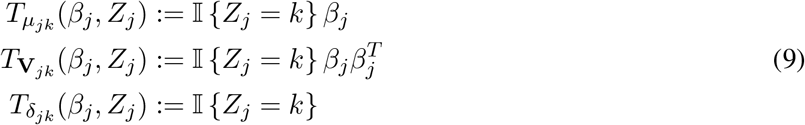

and remains conjugate to the multivariate normal likelihood.

Exponential families play a special role in VI. In particular, that our variational family forms a conjugate exponential family in turn allows us to derive simple coordinate-wise parameter updates using the results of [6]. In particular, letting *q*_−*j*_ be Π_*j*′≠*j*_*q*_*j*′_(*β*_*j*′_, *Z*_*j*′_), we have that for fixed *q*_−*j*_ the ELBo is optimized with respect be to *q*_*j*_ at

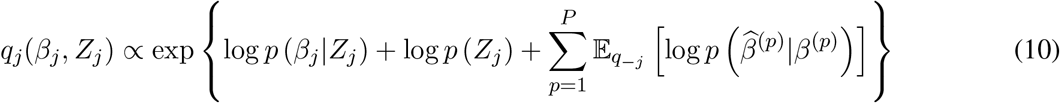

and this is in the same exponential family as the prior. We therefore just need to find the coefficients of the sufficient statistics (Equations 9) in Equation 10, which will give us the optimal values for the natural parameters. We can then solve Equations 8 to obtain the more standard parameters of a mixture of multivariate Gaussians from the natural parameters.

Expanding the exponent of Equation 10 we see (where we abuse notation and use to denote that we are now dropping *additive* constants here):

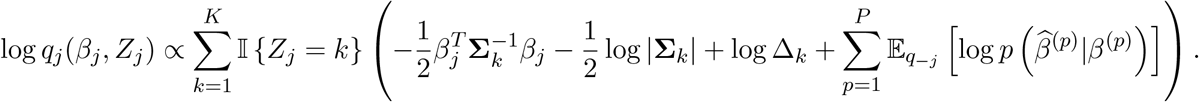

To tackle the terms like 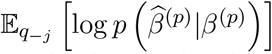 we will drop the (*p*) notation below for convenience, and re-add it once we again consider the likelihood in multiple cohorts. As discussed in Section 4.1.3 we use a low-rank approximation to the LD matrix **X**, and as such throughout we will abuse notation and write **X**^−1^ for the pseudo-inverse of **X**. Importantly, **XX**^−1^ is *not* the identity matrix. As such we write **X**^*°*^ := **XX**^−1^ = **X**^−1^**X**.

Noting that we only care about terms that vary with *β*_*j*_:

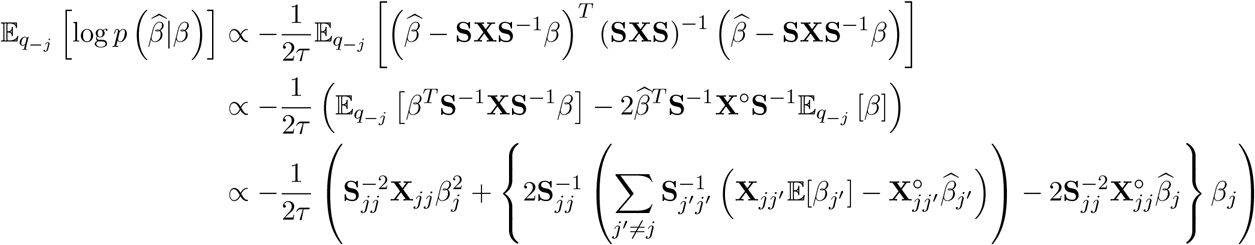

and we can compute these expectations as:

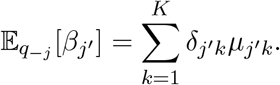

Plugging these into Equation 10, we obtain that the coefficients of the sufficients statistics are

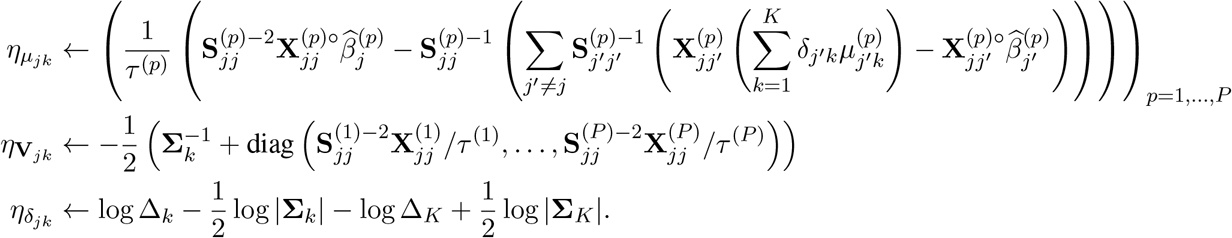

We can then solve Equations 8 to obtain the standard parameterization of our distribution:

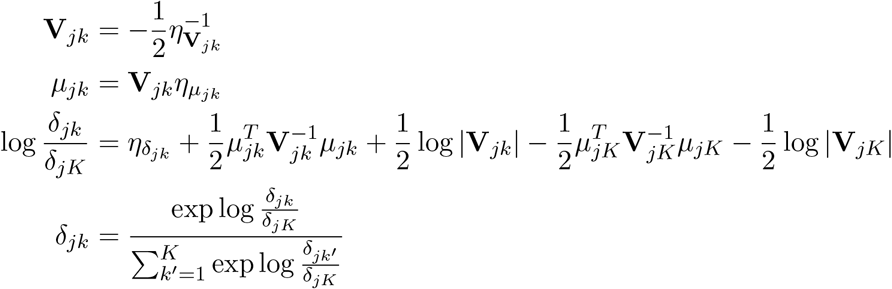

One thing that we can immediately note is that 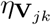 and 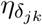 do not depend on the data, 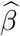, or on the variational parameters at any other position *j*^*′*^. As such, for fixed *τ* and Δ we can immediately set those parameters to their optimal values for all *j* and all *k*. All that remains is finding the optimal 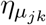 for all *k* and *j*. One option would be to do coordinate ascent, but to take advantage of parallelism, we instead use the fact that the update for 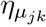 can be viewed as a step in the direction of the natural gradient [1, 6]. As such, call the update for 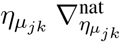, we can then collect this across all *j* and *k* to obtain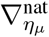. We can then consider a step in the direction of the natural gradient as:

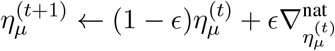

where we use (*t*) to index the gradient step iteration, *ε* to denote a step-size between 0 and 1, and use *η*_*µ*_ to denote the 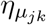 collected across all *j* and *k*. In practice we perform a line-search on *ε* to ensure that the ELBo actually increases after a given step, and then we perform natural gradient steps until the ELBo improves less than a given tolerance threshold.

We now know how to update the variational posterior, but we still need to optimize the hyperparameters *τ* ^(1)^, …, *τ* ^(*P*)^ and Δ_1_, …, Δ_*K*_. Ideally, we would set them by maximizing the marginal likelihood, but that is intractable. Instead, we follow the usual approach of maximizing a lower bound on the marginal likelihood (i.e., the ELBo) with respect to these hyperparameters. We can write the ELBo and note explicitly where the hyper parameters appear

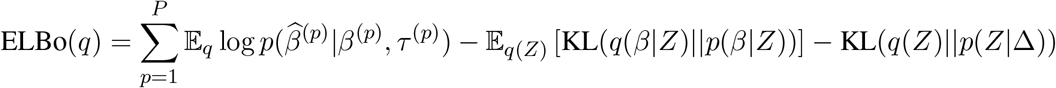

Taking the partial derivative with respect to *τ* ^(*p*)^ we see

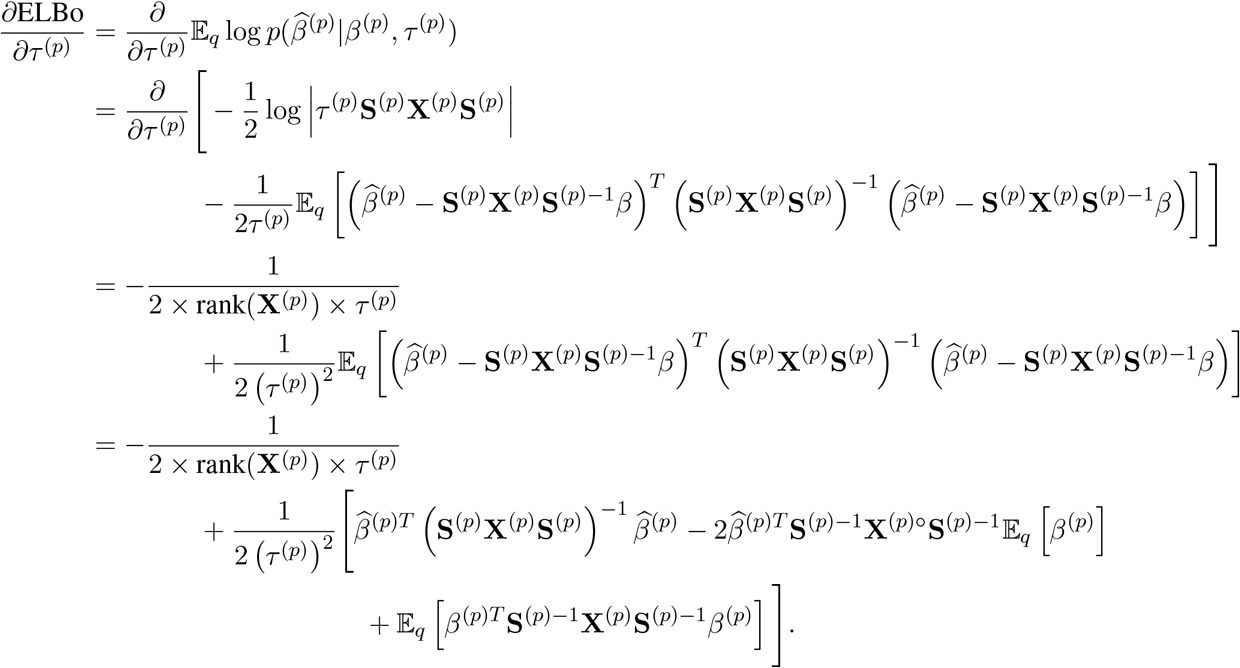

Therefore, the optimal *τ* ^(*p*)^ is

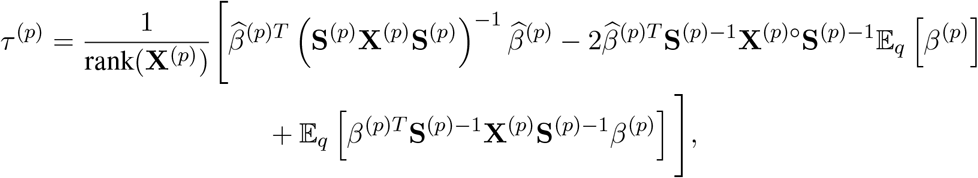

where

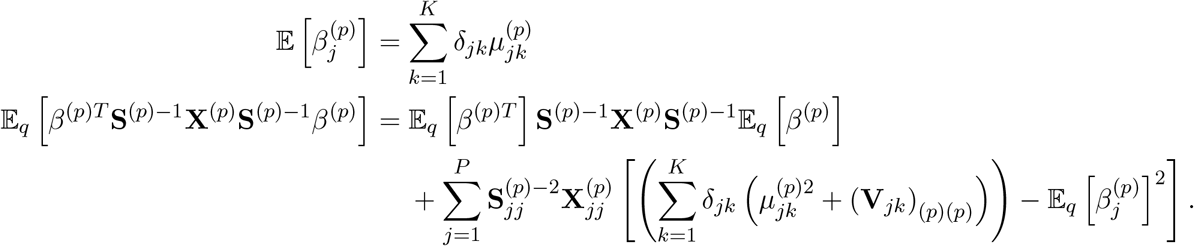

Similarly, taking the gradient with respect to Δ and including a Lagrange multiplier to enforce that 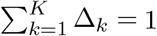, we obtain

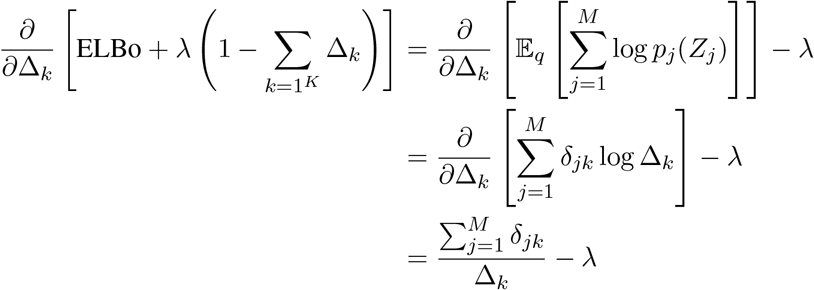

This immediately implies that the optimal Δ_*k*_ is

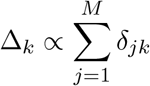

and in fact the constant of proportionality can be obtained by summing across *k*:

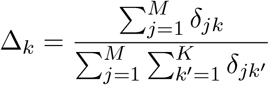

In our implementation we alternately update *q, τ* and Δ until the ELBo stops improving by a sufficient amount, the posterior means remain essentially unchanged, or a user-specified number of iterations is performed.

